# Development differentially sculpts receptive fields across human visual cortex

**DOI:** 10.1101/199901

**Authors:** Jesse Gomez, Vaidehi Natu, Brianna Jeska, Michael Barnett, Kalanit Grill-Spector

## Abstract

Receptive fields (RFs) processing information in restricted parts of the visual field are a key property of neurons in the visual system. However, how RFs develop in humans is unknown. Using fMRI and population receptive field (pRF) modeling in children and adults, we determined where and how pRFs develop across the ventral visual stream. We find that pRF properties in visual field maps, V1 through VO1, are adult-like by age 5. However, pRF properties in face- and word-selective regions develop into adulthood, increasing the foveal representation and the visual field coverage for faces in the right hemisphere and words in the left hemisphere. Eye-tracking indicates that pRF changes are related to changing fixation patterns on words and faces across development. These findings suggest a link between viewing behavior of faces and words and the differential development of pRFs across visual cortex, potentially due to competition on foveal coverage.

---

The receptive field, the portion of visual space from which information is processed, is a fundamental characteristic of the visual system. Receptive fields are found from the earliest stages of the visual system in retinal ganglion neurons^1^, to V1^2^, to high-level visual regions^3–6^ including regions involved in face^4,5^ and word processing^6^. Given behavioral differences across children and adults in both low-level (e.g. visual acuity^7^) and high-level (e.g. face recognition^8^) visual behaviors reliant on receptive fields, it is possible that receptive fields continue to develop across the entire ventral stream after age 5. However, fundamental questions remain unanswered: (1) Do receptive fields in human visual cortex develop during childhood? (2) If so, what is the nature of the development? (3) What is the relationship between receptive field development and viewing experience? Understanding receptive field development will provide fundamental insight into the most basic computation underlying the function of over 30% of the human brain. With disorders such as dyslexia and autism having been associated with atypical brain processing as well as uncharacteristic fixations patterns^9,10^, understanding the link between receptive field development and viewing experience has broad implications in neuroscience.

High-level visual abilities such as reading and face recognition rely on a series of visual computations across the ventral visual stream^11^: a hierarchy of visual areas beginning with V1 and culminating in ventral temporal cortex (VTC) where face^12^- and word-selective^13^ regions supporting face^14^ and word-form perception^15^, respectively, are located. Since neurons across the entire ventral visual hierarchy have receptive fields^3–6,16^ and neurons with similar receptive fields are spatially clustered^2^, the population receptive field (pRF) of neurons in each fMRI voxel can be reliably measured^17^. In each of early (V1-V3) and intermediate visual areas (V4-VO1) in the ventral stream, pRFs systematically tile the visual field and are organized topographically across the cortical surface into visual field maps^16,17^. In high-level ventral regions that are involved in reading^6^ and face recognition^4,5^, pRFs are large and cover the central visual field, generating an over-representation of the fovea, referred to as a foveal bias^18^, rather than a uniform coverage of the visual field.

Theoretical models make different predictions with regards to the first question: *Do pRFs develop across the ventral stream?* One possibility is that pRFs across the entire visual stream are early-developed or innate. This hypothesis emerges from research showing that the wiring of the visual system which determines neurons’ receptive fields and topographic organization is laid out during embryonic development by molecules that guide axon generation and synaptic formation^19–21^. A second possibility is that there is a gradient of development, whereby earlier visual areas develop prior to higher-level regions in the ventral stream. This hypothesis is predicted by empirical findings showing that functional^22–26^ and anatomical^27,28^ development of face and character-selective regions is protracted compared to earlier regions^29^. A third possibility is that pRFs across the entire ventral stream continue to develop during childhood. This hypothesis is suggested by data illustrating that coarse receptive field properties are instilled via embryonic wiring, but that visual experience is necessary to fine-tune them^19–21^, as molecular cues alone cannot specify the precision of adult receptive fields and visual field maps^19,21^.

A second, related question is: *What neural changes occur during development?* One possibility is that development of pRFs and visual maps is associated with qualitative changes from childhood to adulthood. For example, perhaps not all visual field maps beyond V1 are fully formed in children. A second possibility is that developmental changes are quantitative, but not qualitative. This possibility predicts a similar functional organization of visual field maps in children and adults, even as pRF properties continue to be fine-tuned throughout development. An influential theory – eccentricity bias^18,30,31^ – makes specific predictions regarding face- and character-selective regions. In brief, the eccentricity bias theory suggests that because face and word processing require high visual acuity enabled by foveal vision, foveation on faces and words during development leads to the emergence of face- and character-selective regions on existing cortical foveal representations. One version of this theory further suggests that competition between representation of faces and words on foveal resources during development together with left lateralization of the language system in the brain is what generates the adult left brain lateralization for words and right brain lateralization for faces^30,31^. However, the eccentricity bias theory does not make specific predictions regarding the development of pRFs and foveal bias in face- and character-selective regions. One possibility is that pRFs and foveal bias are innate or early-developed, which sculpts the later development of face- and character-selectivity as reported by prior studies^22,25,31,32^. An alternative possibility is that the foveal bias continues to develop throughout childhood, increasing in the left hemisphere within character-selective regions and in the right hemisphere in face-selective regions, consequently enabling more proficient processing of words and faces, respectively.

*Is viewing behavior linked to pRF and visual field map development?* A large body of behavioral research has shown that fixation patterns in adults are task-dependent, placing their foveal resources on task-relevant information. For example, during face recognition, adults tend to fixate on the center of faces^18,33^ (nose bridge) putting informative features^34,35^ at the region with the highest acuity. However, it is unknown if children fixate on faces and words in the same way as adults, or if their viewing patterns develop together with the development of pRFs. If fixation patterns are adult-like in children, even as pRFs develop, it would provide evidence supporting the hypothesis that viewing experience shapes pRFs. However, if fixation patterns change together with pRF development, it would suggest that there is a developmental interplay between pRF formation and viewing experience. In turn, this predicts that in order to scan faces and words like adults, pRFs need to be fully developed.

To address these key questions and elucidate the development of pRFs and visual field maps in the ventral visual stream, we modeled pRFs with functional magnetic resonance imaging (fMRI, see Online Methods) in children (n=26, 5 to 12 years old) and adults (n=26, 22-27 years old). Subjects were scanned as they viewed a sweeping checkerboard bar while fixating on a central stimulus and performing a color-change task. We modeled the pRF of each voxel in the ventral stream as a 2-dimensional Gaussian with a nonlinearity, referred to as compressive spatial summation^4,36^ (CSS). CSS improves pRF fits in higher-level visual areas^4,36^. We examined: (i) if there are qualitative differences across age groups in pRF properties and visual field maps, (ii) if there are quantitative differences across age-groups in pRF size, pRF eccentricity, and visual field coverage (VFC) obtained by the collection of pRFs spanning each visual area, and (iii) if developmental effects differ across ventral visual stream regions.

To examine if there is a relationship between pRF development and viewing behavior, a subset of subjects participated in a behavioral experiment outside the scanner on a different day. Here, participants viewed images of faces and words during a recognition task while their fixations were recorded by an eye-tracker. We tested if fixation patterns on faces and words differed between children and adults and if so, whether they were associated with pRF properties measured separately during fMRI.

## RESULTS

### Early and intermediate ventral visual areas are developed by age 5

All subjects completed pRF mapping. There were no significant differences across age groups in (i) motion during fMRI (adult motion average: 0.7mm±0.33mm, child: 0.89mm±0.2mm; t(39)=1.4, *n.s.*), (ii) fixation behavior during fMRI (t(30)=1.73, *n.s.* **Fig S1A,B**), or (iii) task performance during fMRI (t(14)=1.28, *n.s.*, **Fig S1C**). To test the goodness-of-fit of the pRF model, we measured the mean variance explained by the model for V1 voxels in each participant and compared across age groups. We matched age groups on the variance explained by the pRF model in V1 voxels by excluding 8 children with the lowest V1 model fits and 3 adults with the highest V1 model fits. This matching resulted in no significant differences across age groups in the percentage variance explained by the pRF model across ventral visual regions (**Fig S2C**). These quality assurance metrics ensure that any developmental effects that we may find are not due to differences between age-groups in motion, performance during fMRI, pRF model fits, or measurement noise.

**Figure 1:**
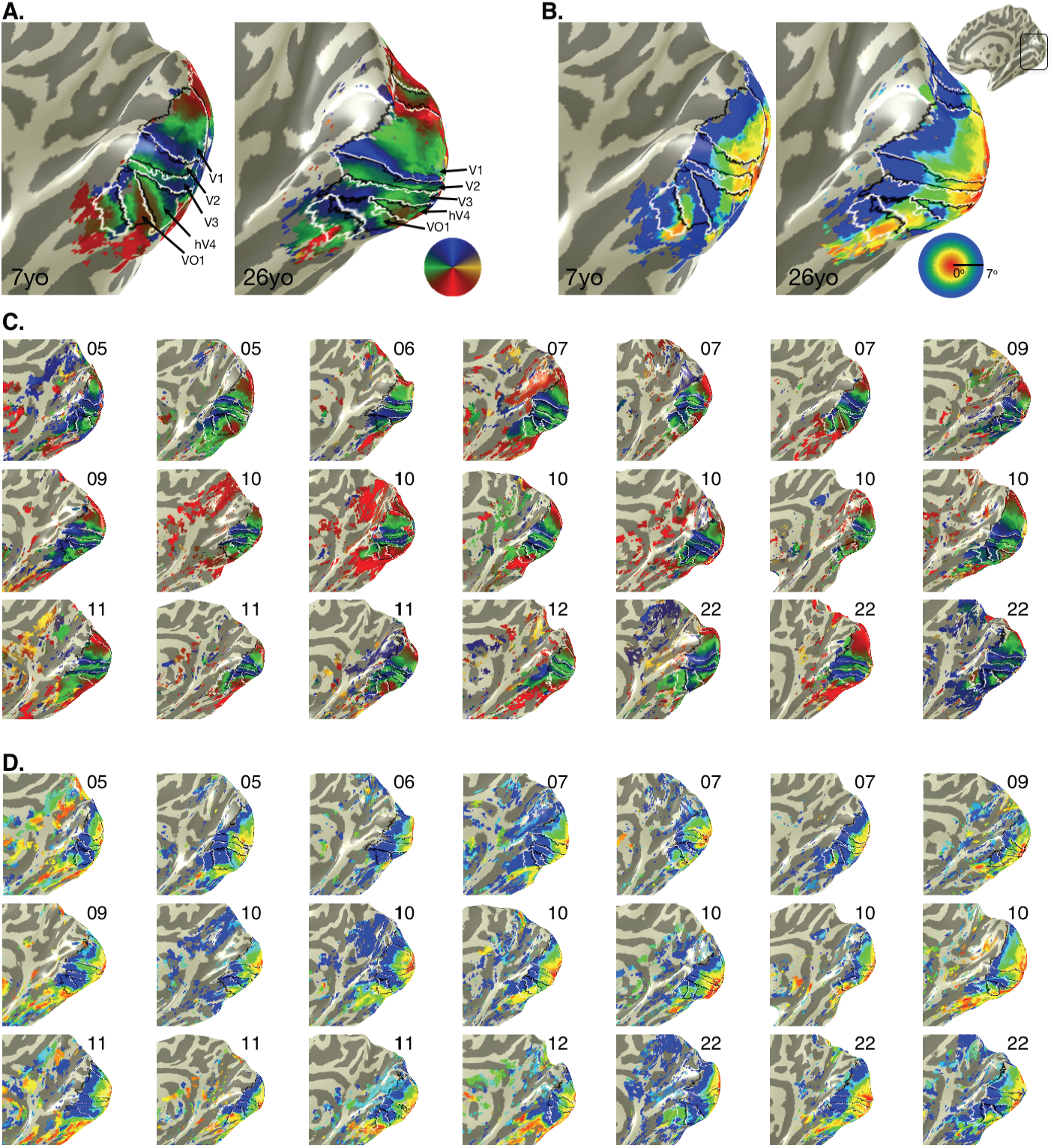
Polar angle and eccentricity maps are qualitatively similar in children and adults. (A) Polar angle maps from an example 7-year-old and 26-year-old. *Color wheel*: polar angle. (B) Eccentricity maps in the same subjects. *Color wheel:* eccentricity. Inset brain shows zoomed region in black outline. (C) Polar angle maps and (D) Eccentricity maps in the right hemisphere of all child participants and 3 example adults. Numbers indicate the age of the participant. Maps are of the central 7° and are thresholded at 5% variance explained, voxel level*. Lines:* boundaries of visual field maps. Polar angle and eccentricity maps of all subjects and both hemispheres are shown in Figs S3-S6.

Examination of the topographic organization of polar angle and eccentricity maps revealed that these maps were qualitatively similar across age groups (**Fig 1, Figs S3-S6**, all participants’ maps). That is, children, like adults, displayed (i) a series of mirror-reversed polar angle maps (**Fig 1A,C**) emerging from a hemi-field representation in and around the calcarine sulcus (corresponding to V1) and (ii) two sets of large-scale eccentricity maps, one spanning the occipital cortex, in which eccentricities progressively increase from posterior to anterior, and one in VTC, in which eccentricities progressively increase from lateral to medial (**Fig 1B,D**).

Using polar angle and eccentricity maps, we successfully defined visual areas V1 through VO1 bilaterally in all 18 children and all 23 adults (**Figs 1, S3-S6**). The cortical volume of visual field maps was slightly (<5%) smaller in children than adults (**Fig S2A**), but like adults, over 90% of voxels were driven by the mapping stimulus and could be modeled by a pRF (**Fig S2B**).

Notably, there were no significant differences across age groups in mean pRF size (**Fig 2A**) or mean pRF eccentricity in V1–VO1 (**Fig 2B**) (*Fs*(1,195) < 1.1, *Ps* > 0.36). Furthermore, in V1-VO1, there was no correlation between mean pRF size and age (-0.11<*Rs*(41)<0.12, n.s) or mean eccentricity and age (-0.04<*Rs*(41)<0.29, n.s). In children’s V1–VO1, like in adults’, pRF size linearly increased with eccentricity (**Fig 2C**). Likewise, there were no significant differences across children and adults in either the slopes (*F*_(1,195)_=0.39, *n.s.,* 2-way ANOVA with factors of visual area and age group) or intercepts (*F*_(1,195)_=2.98, *n.s*., 2-way ANOVA) of the pRF size vs. eccentricity line fits in V1–VO1. In children, like adults, pRF size also increased across the visual hierarchy, demonstrated by the progressive steepening of slopes of the size vs. eccentricity line from V1 to VO1 (**Fig 2C, Fig S7**) and the systematic increase in mean pRF size ascending the visual hierarchy (**Fig 2A**).

As there were no quantitative differences in pRFs properties across children and adults in V1-VO1, the visual field coverage (VFC) obtained by the collection of pRFs spanning each of these visual field maps was strikingly similar across children and adults (**Fig 2D**). In each of V1 through VO1, the VFC was largely uniform and spanned a hemi-field in each hemisphere in both children and adults. There was no significant difference in the VFC of V1 through VO1 across development (main effect of age group: F_(1,244)_=1.76, *n.s.*). Together, these analyses reveal that past the age of 5, children have adult-like polar angle and eccentricity maps, and adult-like pRF properties and VFC in V1–VO1.

**Figure 2:**
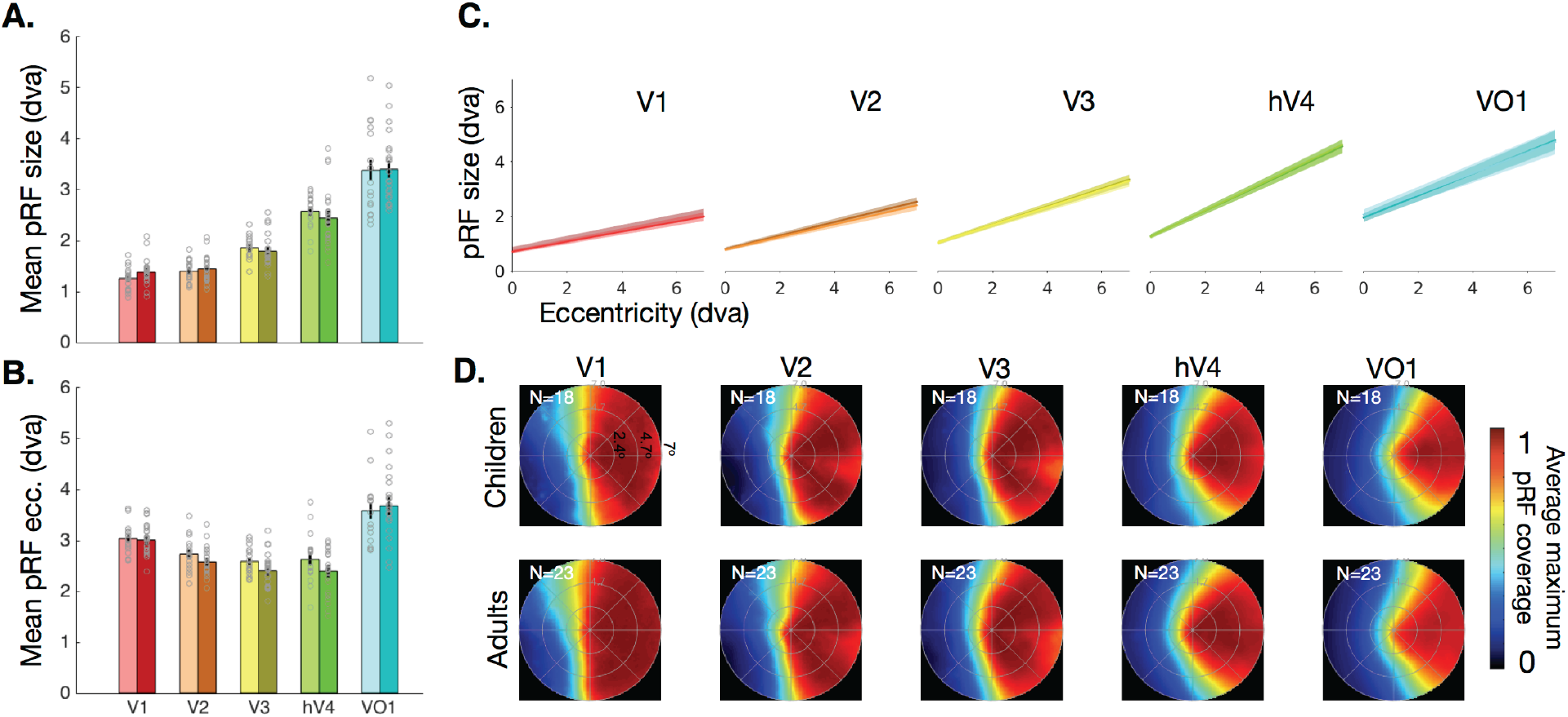
pRF and visual field coverage in V1-VO1 are quantitatively similar across children and adults. (A) Mean pRF size in V1-VO1 across 18 children (light colors) and 23 adults (dark colors); *Errorbars:* standard error*. Gray circles:* individual subject data. Each circle is a participant. (B) Mean pRF eccentricity in V1-VO1 of the same subjects. *Gray circles:* individual subject data. (C) pRF size vs. eccentricity relationship is similar across age groups. The line of best fit (solid line) and the standard error (shaded region) illustrates the relationship between pRF eccentricity and size in units of degrees of visual angle (dva). Adults are shown in dark colors (n=23), children (n=18) in light colors (each age group is shown separately in Fig S7). Fits are calculated in each subject, slopes and intercepts are then averaged across subjects. (D) Visual field coverage of V1-VO1 computed using the average maximum pRF density coverage for each subject and then averaged across subjects. Maps are averaged across hemispheres by flipping the right hemisphere data*. Top:* children. *Bottom:* adults. Number of participants is indicated in the top-left of each panel Inner to outermost ring segments correspond to 2.4, 4.7, and 7 degrees of visual angle (dva).

### The VFC of face- and word-selective regions develops after age 5

To examine if pRFs in high-level regions develop with age, we next defined face- and word-selective regions in all subjects using an independent localizer experiment (**Fig 3A**, Online Methods), and compared across age groups mean pRF size, mean pRF eccentricity, and the VFC of each of these regions. We focus on face-selective responses on the posterior fusiform gyrus (pFus-faces) and word-selective responses in the posterior occipitotemporal sulcus (pOTS-chars; Online Methods) because (i) these regions are proximal to the VO1/VO2 transition in VTC, and (ii) a substantial number of voxels in these regions were modulated by the checkerboard mapping stimulus and therefore could be fit by the pRF model (**Fig S2E,F**). It is noteworthy that in face-selective pFus-faces and word-selective pOTS-chars children had (i) significantly more voxels that were modulated by the pRF mapping stimulus than adults (**Fig S2E,** F(1,115) = 5.68, *P*<0.02) and (ii) significantly higher percentage variance explained by the pRF model compared to adults (**Fig S2F**, *F*_(1,114)_ = 8.24, *P*<0.005, 2-way ANOVA with factors of ROI and age). In general, the size of these regions was not significantly different across age groups (**Fig S2D**, *F*_(1,115_) = 0.44,*n.s.*), except that pFus-faces was numerically larger in adults than children. This difference in voxel number is even smaller considering children had more voxels driven by the bar stimulus in face-selective regions than adults. Additionally, in these regions, there was no correlation between mean pRF size and age or mean eccentricity and age either when considering all subjects or just children (0.35 > *Rs* > −0.24, *n.s.*), justifying the grouping of children into one group.

**Figure 3:**
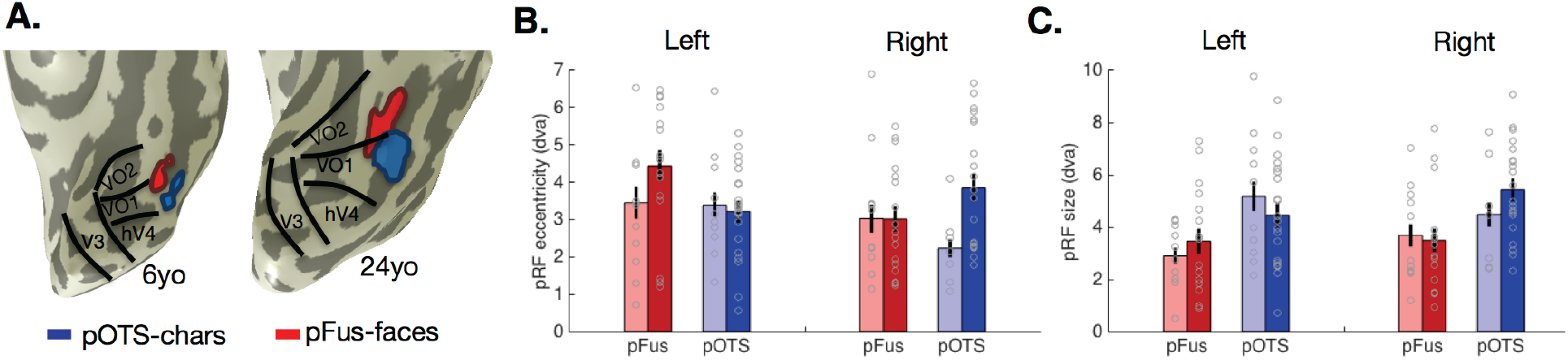
Mean pRF eccentricity and size in pFus-faces and pOTS-chars change with development. (A) pFus-faces (red) and pOTS-chars (blue) on the left ventral temporal lobe in representative child and adult participants. Black lines: boundaries of visual field maps. (B) Mean pRF eccentricity in pFus-faces (red) and pOTS-chars (blue) in children and adults, units are degrees of visual angle. Children are in light colors. (C) Mean pRF size in pFus-faces (red) and pOTS-chars (blue) in children and adults, units are degrees of visual angle. (B-C) Error bars: standard error across subjects of an age group. *Gray circles:* individual subject data. Left pOTS-chars: 12 children, 22 adults; Right pOTS-chars: 8 children, 21 adults. Left pFus-faces: 11 children, 18 adults; Right pFus-faces 14 children, 18 adults.

In opposition to preceding visual field maps, we found development of pRF properties in pFus-faces and pOTS-chars that varied across hemispheres and regions. Specifically, pRF centers become more eccentric in the left hemisphere for pFus-faces and more eccentric in right hemisphere for pOTS-chars (**Fig 3B**). A 3-way ANOVA on pRF eccentricity with factors of age, hemisphere, and ROI revealed a significant three-way interaction (F_(1,111)_=4.33; p = 0.03) and a significant effect of age (F_(1,111)_ = 4.83, p = 0.03) A separate 3-way ANOVA on pRF size with the same factors reveals a significant effect of ROI (F_(1,111)_ = 13.99, p = 0.0003), with pRFs sizes in pOTS-chars about 56% larger than in pFus-faces, and a trending but non-significant three-way interaction (p=0.1; **Fig 3C**).

**Figure 4:**
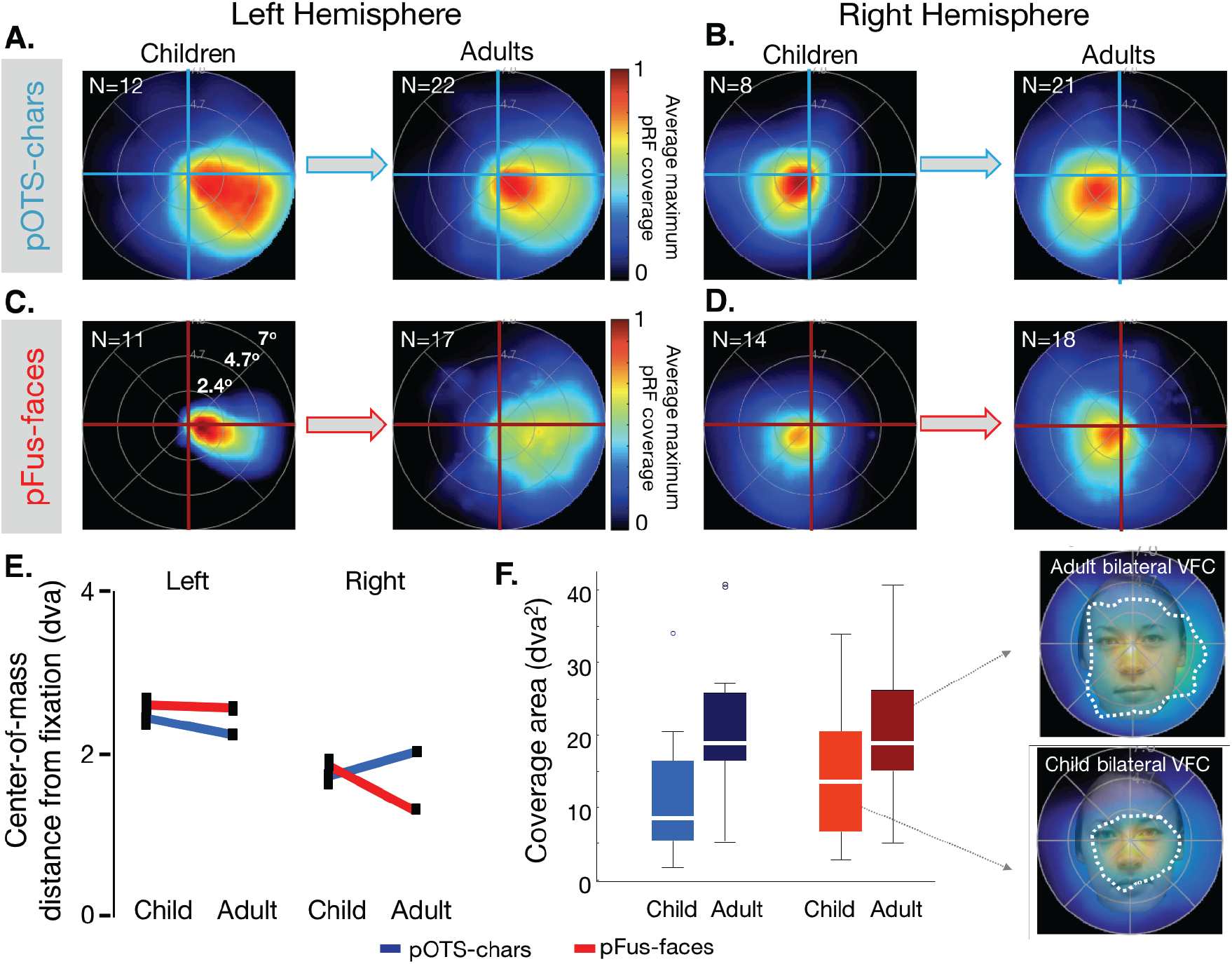
Hemispheric differences in pRF coverage emerge across development in face- and character-selective cortex. (A-D) For each region, visual field coverage (VFC) was calculated using the average maximum pRF density coverage for each subject and then averaged across subjects. The number of subjects used to produce the VFC is indicated in the upper left of each panel. Innermost to outermost rings correspond to 2.4, 4.7, and 7 degrees of visual angle (dva), respectively. Arrows illustrates development from childhood to adulthood. (A) VFC of left pOTS-face. (B) VFC of right pOTS-chars. (C) VFC of left pFus-faces. (D) VFC of right pFus-faces. (E) Center-of-mass (CoM) distance of the VFC from fixation illustrated for left and right pOTS-chars and pFus-faces across children and adults. A CoM distance of 2 indicates that the VFC is 2 degrees of visual angle from fixation. *Errorbars:* jackknife standard errors. (F) *Left:* VFC of bilateral pFus-faces and pOTS-chars in children and adults; *white*: indicating the median; *box:* 25^th^ and 75^th^ percentiles; *whiskers:* range. *Right:* overlay of the VFC of bilateral pFus-faces in adults (top) and children (bottom) on a face about 1 m from the observer (corresponding to ∼6.5 dva). *Dashed white contour:* 50% density contour of the VFC. This contour covers more of the average-sized face in typical viewing distance in adults than children.

As the visual field coverage obtained by the collection of pRFs spanning a region depends on the distribution of pRF sizes and eccentricities, subtle development in mean properties may have a profound effect on visual field coverage (VFC) in face- and word-selective regions. To examine this possibility, we estimated the VFC of face- and word-selective regions in each subject, separately for each hemisphere, and then measured the mean VFC of these regions across participants of an age group. As in V1-VO1 there were qualitative similarities in the VFC of face- and word-selective regions across age groups. In both children and adults, the VFC of these regions exhibited a contralateral preference, a foveal bias, and a greater coverage of the lower than upper visual field (**Fig 4A-D**), as reported previously in adults^4,6^. That is, in each hemisphere, pRFs of face and word-selective regions covered more prominently the contralateral and central visual field than the ipsilateral or peripheral visual field

pFus-faces and pOTS-chars, however, differ in their developmental patterns across hemispheres. Specifically, we find significant changes in the VFC obtained by pRFs of pFus-faces and pOTS-chars across children and adults (2-dimensional Komolgorov-Smirnov (K-S) test comparing the VFC of children and adults: left pOTS-chars, K-S = 0.31, p<0.01; right pOTS-chars: K-S = 0.25, P<0.01) and pFus-faces (right pFus-faces, K-S = 0.13, p<0.01; left pFus-faces K-S = 0.32, p<0.01).

To quantify these developmental changes in the VFC, we computed the center-of-mass of the VFC (CoM, reflecting how far the center of the VFC is from fixation, see Online Methods) in each region and age group. In the left hemisphere, the CoM shifts towards the fovea across development in left pOTS-chars (**Fig 4A, 4E-left**), becoming in adulthood closer to the fovea compared to neighboring left pFus-faces (**Fig 4C, 4E-left**). In the right hemisphere, developmental changes in VFC are reversed: in the right pOTS-chars pRFs shift away from the fovea (**Fig 4C, 4E-right**), while in right pFus-faces the CoM moves towards the fovea (**Fig 4D, 4E-right**). Despite no significant difference in ROI size between groups (**Fig S2D**), adult pFus-faces are ∼30% larger than children. To test if ROI size influences developmental results, we dilated children’s pFus-faces to match the mean adult size, and repeated these analyses. Results remain the same (**Fig S8**), verifying that between-group differences stem from pRF development.

In addition to developmental changes in the CoM, we find significant developmental increases in the total extent of the VFC. That is, the total area of the VFC spanned by pRFs across bilateral pFus-faces and bilateral pOTS-chars significantly increases by ∼7 square degrees of visual angle from childhood to adulthood (main effect of age, F_(1,86)_=5.64, p<0.03, **Fig 4F**). Together, these data reveal differential development of the VFC in face- and word-selective regions across hemispheres, and an increase in the total extent of VFC.

### Development of viewing patterns mirrors pRF changes in high-level visual regions

Previous work suggests that optimal viewing behavior involves central fixations, as the center of the stimulus is the most informative region for recognition of faces and words. This framework predicts similar fixation in children and adults. However, our finding of development of the VFC in face- and word-selective regions may impact viewing behavior on faces and words, respectively. We hypothesized that if pRF coverage guides natural viewing behavior, the optimal behavior would be to place the VFC, not the fovea, onto the center of stimuli. For children, this predicts fixations that are biased off of the center, resulting in systematic shifts in the viewing of faces and words across children and adults. Specifically, the neural data make three predictions: (i) due to the larger foveal bias in adults, they will show more central fixations than children, (ii) if the VFC in right pFus-faces guides fixation, children’s fixations on faces will be more rightward and upward biased than adults, and (iii) if left pOTS-words drives fixations on words, children’s fixations on words will be more leftward and upward biased than adults.

We assessed natural viewing of faces and words in a subset of our participants (12 children and 11 adults) in a separate behavioral experiment. Outside the scanner, each participant first viewed a series of images from different categories (including faces and pseudowords) and performed a one-back task. Then, participants completed a surprise, self-paced old-new recognition task during which their eye movements were recorded with an eye-tracker (Online Methods). We then determined if free-viewing fixation patterns followed the predictions of the visual field coverages obtained from the fMRI experiment inside the scanner while participants were fixating.

**Figure 5:**
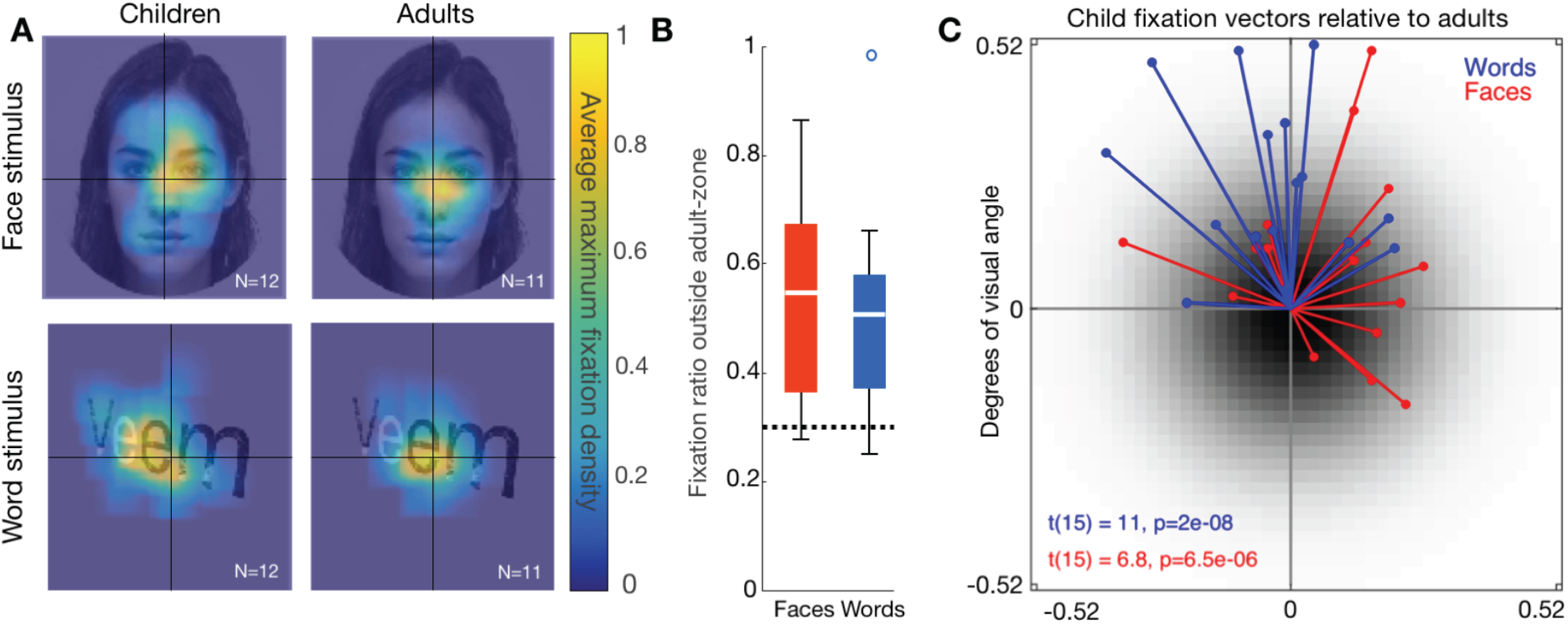
Fixation patterns on face and word stimuli become more central with development. (A) Average maximum fixation density maps produced from 12 children and 11 adults during a free-viewing recognition task are overlaid on an example face (top row) and pseudoword (bottom row) stimulus. Fixation density patterns in adults are more centrally placed on faces and words than in children. (B) The mean ratio of fixations in children made outside of an adult-like zone defined as the region where 70% of adults or greater fixated on the stimulus. The dashed line at 0.3 denotes the expected adult value. *White:* median; *Box:* 25^th^ and 75^th^ percentiles; *Whiskers:* range; *Circles:* outliers; *C:* children; *A:* adults; *red:* unfamiliar faces; *blue:* pseudowords. (C) Vectors describing the bias in child fixation densities for all face (red) and pseudoword (blue) stimuli relative to the center of adult fixation densities for each stimulus. Each vector is the bias for a particular stimulus. Black Gaussian center represents the centrally-biased adult fixation densities.

Results show that fixation locations on both face and pseudoword stimuli differed between children and adults. As shown for the example stimuli, adults foveate more centrally within face and pseudoword stimuli, while children fixations are more eccentric across the stimulus expanse (**Fig 5A**). To quantify differences in fixation patterns across age groups, we measured the region of the image in which adults make most of their fixations by calculating for each face and pseudoword stimulus the central region in which adults made 70% of their fixations. Then, we calculated for each child and each image the proportion of fixations made outside of the adult fixation zone and then derived the mean proportion of such fixations across child participants. Results indicate that children fixate significantly outside of the central adult fixation zone for both face (t(11)=4, p<0.01) and word (t(11)=3.63, p<0.01) stimuli (**Fig 5B**), whereby about 50% of their fixations are outside the adult central fixation zone, even as they make fewer fixations than adults (**Fig S9**).

Critically, it is not the case that children make more variable fixations than adults, as they show systematic biases in their fixation patterns. Notably, these biases mirror the asymmetries in visual field coverage of face- and word-selective regions in their dominant hemispheres. As shown for the example stimuli, children tend to bias their fixations towards the upper right side of faces (**Fig 5A**) which puts the VFC of right pFus-faces, which is biased to the left and lower visual field, in a location where it optimally covers the face. Similarly, children tend to fixate on the leftward aspect of words (**Fig 5A**), putting the VFC of the left pOTS-chars, which covers the right horizontal visual field, in a place where it optimally covers the word. We quantified this fixation bias by calculating the center-of-mass of fixation densities on each face and pseudoword stimulus separately for adults and children. In **Fig 5C**, we plot for each image the vector representing the displacement of child fixation densities relative to adults. Strikingly, children are significantly biased to the upper right quadrant for faces (t(15)=6.8, p<0.001) and the upper left quadrant for words (t(15)=11,p<0.001, Online Methods). Importantly, there is no stimulus on which children fixate into the visual field quadrant containing their pRFs (lower left for faces, lower right for words), which would move their VFC further from the stimulus. Together, behavioral measurements during natural viewing strikingly show that both adults and children fixate in a manner that puts their visual field coverage in face- and word-selective regions on the informative region in the visual stimulus.

## DISCUSSION

Modeling population receptive fields in human visual cortex for the first time in children, we find evidence for differential trajectories of development within the ventral stream and across hemispheres. Early and intermediate visual areas V1–VO1 are developed early, while high-level visual regions in VTC show protracted development in representation of the fovea and visual field coverage from childhood to adulthood. Importantly, fixation patterns on face and pseudoword stimuli during natural viewing demonstrate a link between viewing behavior and developmental changes in the visual field coverage by pRFs in face- and word-selective regions. These data provide insight into the possible role of visual experience in sculpting the spatial window through which high-level visual regions process visual information.

We find no qualitative or quantitative difference in visual field topography or pRFs in early and intermediate visual areas V1-VO1 across children and adults. These data suggest that receptive field properties and visual field maps in the human ventral stream are developed by age 5, consistent with predictions of developmental theories based on research in animal V1^19–21^ and retinotopic mapping in V1-V3^29^. Whether pRF development in visual field maps of the dorsal stream^37^ follows a similar developmental time course is a topic for future research. By contrast, we find developmental changes in pRF centers and visual field coverage in face- and word-selective regions beyond age 5 in the same participants. Thus, our results provide the first evidence that development of V1-VO1 precede that of downstream ventral regions. These findings hold important implications for understanding the origins of functional architecture in the ventral stream for two reasons. First, they suggest that the early development of visual field maps V1-VO1 may be the neural scaffold that constrains the later emergence and ultimate topography of neighboring high-level visual regions^18^. Second, our findings that pRFs develop beyond V1 and after age 5 drastically extend both the length of time and expanse of visual cortex where pRF development occurs compared to what is known from research in neonate animal V1^19–21^. Future research using participants spanning a broader age range as well as longitudinal measurements can elucidate the developmental trajectory of pRFs in visual cortex across childhood.

The eccentricity bias theory^18,30,31^ suggests that foveation on faces and words during natural viewing anchors the processing of these stimuli to regions in VTC representing the fovea. Consistent with this view, in both children and adults, the visual field coverage in face- and word-selective regions is foveally biased, providing a more substantial coverage of the central than peripheral visual field. Future longitudinal research in younger participants will determine whether the over-representation of the central visual field emerges before or together with selectivity to faces or words.

Unpredicted by the eccentricity bias theory, our data show that spatial representations in these high-level regions continue to develop from childhood to adulthood. In fact, both the foveal bias and the overall visual field coverage obtained by pRFs in face and word-selective regions increase from childhood to adulthood. These findings argue against the hypothesis that face or word selectivity develop on top of a mature foveal bias and spatial representation. The expansion of the visual field coverage in face- and word-selective cortex and increased foveal bias may involve proliferation of dendritic arbors and synapses to support the increased pooling of information. Thus, pRF development may be associated with microstructural cortical tissue growth that has been observed in face- and word-selective regions^27^.

Notably, the development of pRF properties and visual field coverage also varied by hemisphere across face- and word-selective regions. Word-selective regions became more foveally biased in the left-hemisphere, where previous research has demonstrated lateralization for word-form processing and reading^38–40^. By contrast, face-selective regions became more foveally biased in the right hemisphere where face processing is thought to be lateralized^41,42^. Intriguingly, at the same time, visual field coverage shifted away from the fovea for face and word-selective regions in their non-preferred hemispheres. This pattern of development has important implications for the theory that reading and face recognition compete for foveal representations^31,43^ because it provides striking evidence for a competitive push-pull mechanism in which the foveal over-representation increases in one hemisphere and decreases in the other, in an opposing manner across hemispheres for faces and words. Additionally, the retreat of pRF coverage from the fovea in non-preferred hemispheres mirrors previous observations of development reductions in responses to nonoptimal stimuli^25^.

Critically, developmental increases in both the foveal bias and visual field coverage in face and word-selective regions measured with fMRI during fixation were associated with developmental changes in fixations on faces and words measured during natural viewing outside the scanner. These data not only bridge for the first time development of spatial processing in high-level vision and real-world viewing behavior, but also demonstrate a direct relationship between pRF properties of cortical regions and viewing behavior of complex stimuli. While our research does not inform whether behavioral changes in fixation patterns on face and word stimuli drive the development of pRFs in face and word-selective regions, or if pRF development drives the behavioral changes, we note that both children and adults fixate in a way that places the VFC of face- and word-selective regions in an optimal location to process these stimuli. In children, left pOTS-chars is less foveal, more rightward and lower-field biased compared to adults. Consequently, their fixations on words are more left and upper-field biased than adults. Likewise, in children, right pFus-faces is less foveal and more leftward biased compared to adults. Therefore, children’s fixations on faces are more rightward biased than adults. These results, therefore, suggest a tripartite relationship between development biases in visual field coverage in high-level regions, fixations patterns, and hemispheric lateralization. It is likely an iterative and bidirectional process whereby learning optimal fixation locations (e.g. the center of a face) produces changes in the biases of visual input, altering pRF centers and size to optimally cover regions of visual interest. Future research examining pRF development in readers of languages demanding different fixation patterns on words (e.g. Hebrew or Chinese) may explicate the interplay between viewing behavior and hemispheric lateralization.

Our findings are important not only for elucidating the development of a fundamental computation – spatial processing by receptive fields – in the human ventral stream and showing its relation to viewing patterns, but also for providing an innovative methodology and computational framework for investigating development of computations across cortex more broadly. As receptive fields are a basic hallmark of neurons in sensory cortical systems (e.g. auditory^44,45^ or somatosensory^46^ cortex), as well as characterize complex cognitive tuning (e.g. to numerosity^47,48^) our novel approach can be applied to quantitatively examine development of cortical function throughout the brain. Likewise, our findings lay fundamental groundwork towards understanding abnormal cortical processing as well as potential maldevelopment in atypical populations, including developmental prosopagnosia^49^, dyslexia^10^, and autism^9,50^.

In sum, we find that early-developed visual field maps in the human ventral visual stream may provide a neural scaffold that shapes the organization of high-level visual regions and that the development of pRFs in high-level visual areas involved in face and word processing is linked to changing viewing patterns on faces and words. Together, these data suggest that both the spatial window through which a region of cortex processes information and our visual experience of complex stimuli changes from childhood to adulthood.

## ACKNOWLEDGEMENTS

This research was funded by the NSF Graduate Research Development Program (grant DGE-114747) and Ruth L. Kirschstein National Research Service Award (grant F31EY027201) to JG, and the NIH (grants 1ROI1EY02231801A1, 1RO1EY02391501A1) to KGS and training grant 5T32EY020485 supporting VN.

## DATA AVAILABILITY

All code relevant to data analysis for the main findings (Figures 1-5) will be available on github.com/VPNL upon request. Any source data relevant to these analyses will also be made available upon request. The majority of the code used in this study was derived from scripts and functions available through the open-source vistasoft code library: https://github.com/vistalab/vistasoft.

## ONLINE METHODS

### Participants

26 neurologically typical children ages 5-12 years (mean age 8.5 ± 2.2 years, 12 females) and 26 adults ages 22-28 years old (mean age 24 ± 1.6 years, 9 females) participated in these experiments. Age ranges were chosen in children to *(i)* maximize a wide dynamic range of functional and structural development reported previously^22,24–26^ and *(ii)* maximize the success of MRI measurements without having to discard a substantial number of participants due to excessive motion in the scanner, which is a common issue with pediatric neuroimaging^51^. Because our goal was to link functional and behavioral changes, and our experiments required maintaining central fixation, we could not make measurements on younger children where acquiring such data is unfeasible. A similar range of ages was chosen in adults when most structural and functional development in VTC is thought to be near completion^52,53^. Following data quality thresholds discussed below, 8 children and 3 adults were excluded from further analysis (18 children, 23 adults remain). Participants had normal or corrected-to-normal vision and were screened to have no prior or current psychiatric conditions. All procedures were approved to be in accordance with the Institutional Review Board of Stanford University. Prior to the experiment, adult participants and parents provided written informed consent, and children provided written assent.

Each subject participated in several sessions completed over the course of a few months to distribute measurements and avoid fatigue. Each of the following sessions was thus performed on a different day: (i) Participants under the age of 18 completed training in a mock scanner employing live feedback of head motion during the viewing of a 15-minute movie. This acclimated the participants to the scanner environment and reduced motion. Participants were advanced to functional and anatomical scanning if they could lie still (less than 2.4 mm of head motion) for the duration of mock scanning. (ii) Children completed the recognition memory task with eye tracking outside the mock scanner on the same day in which they participated in training; adults completed this task after scanning was completed. (iii) All participants participated in an MRI session in which we obtained anatomical MRI brain volumes which were used to register data across sessions and obtain cortical surface reconstructions of each brain. (iv) All participants participated in an fMRI session in which we measured brain responses to stimuli of various categories (referred to as localizer experiment). (v) All subjects participated in an fMRI session composed of four runs of pRF mapping.

#### Data acquisition

##### Quantitative magnetic resonance imaging (qMRI)

Quantitative MRI measurements are obtained from the protocols set forth in^54^. T1 relaxation times were measured from four spoiled gradient echo (spoiled-GE) images with flip angles of 4, 10, 20, 30 (TR=14 ms, TE=2.4 ms) and a scan resolution of 0.8 mm × 0.8 mm × 1.0 mm. For the purposes of removing field inhomogeneities, we collected four additional spin echo inversion recovery (SEIR) scans with an echo planar imaging (EPI) read-out, a slab inversion pulse, and spectral spatial fat suppression. The SEIRs were acquired with a TR of 3.0 sec, echo time set to minimum full, and 2x acceleration. The inversion times were 50, 400, 1200, and 2400 ms, and were collected at a 2.0 mm × 2.0 mm in-plane resolution and a slice thickness of 4.0 mm. An artificial T1-weighted anatomy optimized for tissue segmentation was produced for each subject from these quantitative measures which were used for surface reconstruction and visualization of retinotopic data.

##### Functional MRI

Data were collected on a 3-Tesla GE Discovery MR750 scanner (GE Medical Systems) at the Center for Cognitive Neurobiological Imaging at Stanford University using a phase-array 32-channel head coil. Functional data for the category localizer were collected with a simultaneous multi-slice EPI sequence with a multiplexing factor^55^ of 3 to acquire near whole-brain (48 slices) volumes at TR=1s, TE=30ms. Data were acquired at a resolution of 2.4mm isotropic voxels with one-shot T2*-sensitive gradient echo sequence with slices aligned parallel to the parieto-occipital sulcus. Functional data for retinotopic mapping were of similar resolution and orientation but collected on a 16-channel head coil, TR=2s, acceleration factor of 2, 28 slices.

##### fMRI category localizer experiment

The purpose of this experiment was to identify those voxels whose neural response preferred either faces or words in order to localize face- and word-selective cortex as functional regions of interest. During scanning, subjects completed 3 runs, each 318 s long, of an experiment presenting subjects with stimuli from 5 categories each with two subcategories (Faces: child, adult; Bodies: whole, limbs; Places: corridors, houses; Objects: cars, guitars; Characters: words, numbers) as described previously^23,27,56^. Images of a category were presented in 4 s miniblocks at a rate of 2 Hz and did not repeat across miniblocks or runs. Each category was shown 8 times in a run in counterbalanced order interleaved with blanks. Subjects fixated on a central dot and performed an oddball detection task of phase scrambled images.

##### pRF mapping experiment

The purpose of this experiment was to model in every voxel the region of the visual field that is capable of eliciting a response from that voxel, namely its receptive field. Subjects completed 4 runs of an experiment in which subjects fixated on a central stimulus and were required to indicate via a button-press when the central stimulus changed color. Black and white checkerboard bars (width = 2° of visual angle, length = 14°) were swept across the screen during each run which lasted 3 minutes and 24s. Bars swept the visual field in 8 different configurations in each run (4 orientations: 0°, 45°, 90°, 135°, each orientation was swept in 2 directions that were orthogonal to the bar). Same as^17,57^. Eye-tracking and fixation task performance were collected on a subset of children and adults (**Fig S1**). Fixation performance on subjects was tracked with the Eyelink software (http://www.sr-research.com/). Blinks, labeled by the Eyelink software, were removed from the timecourse data of the recorded eye by scrubbing with a 100ms window on either end of the blink. Fixation data was then plotted for each subject. Only subjects that made fewer than three saccades (2° in size) during a mapping run were included for analysis. Due to the scanner environment, size of participants’ head, and time constraints, not all subjects could be eye-tracked during pRF mapping (eye tracking data was obtained for 25 children and 6 adults). Fixation task performance was also only collected on a subset (8 children, 7 adults) of subjects due to button box malfunction. All subjects, however, were trained on proper fixation technique during the recognition memory task (see Measuring Fixation Patterns below), and all subjects included in the analysis that underwent eye-tracking in the scanner fixated successfully, with no difference between age groups. As a reminder, we also observe no difference in pRF properties or pRF model performance in V1 between children and adults, further suggesting proper fixation performance, as improper fixation significantly impacts pRF size estimates^58^.

#### Data Analysis

##### Anatomical data analysis

Both the spoiled-GE and the SEIR scans were processed using the mrQ software package in MATLAB to produce T1-weighted maps^54^. The mrQ analysis pipeline corrects for RF coil bias using SERI-EPI scans, producing accurate proton density (PD) and T1 fits across the brain. The full analysis pipeline and its published description can be found at (https://github.com/mezera/mrQ). An artificial T1-weighted anatomy was produced for each subject from these quantitative measures which were used for surface reconstruction and visualization of retinotopic data. Anatomical images for each subject were segmented through FreeSurfer (https://surfer.nmr.mgh.harvard.edu/), the resultant tissue segmentation was hand-corrected for classification errors. Functional data were restricted to the cortical ribbon by growing a 3-voxel thick (1 mm isotropic voxels) ribbon from the gray-white matter boundary.

##### fMRI data analysis

Data were processed and analyzed in MATLAB using mrVista software (http://github.com/vistalab) as in previous publications^23,27^. Functional data were aligned to the artificial T1-weighted volume. Functional data were unsmoothed, always analyzed within the individual subject native brain anatomy space, and were restricted to the cortical ribbon.

Functional data were motion corrected both within and between scans. Any subjects who moved more than 2 voxels within a scan were either excluded from data analysis or invited back for another session, such that children and adults were matched for data quality as shown in **Fig S2C**. There was no significant difference in motion during scanning between groups (see Results). To ensure there were no group differences between children and adults resulting from differences in data quality, age-groups were matched for the mean percentage variance explained of the pRF model across voxels in V1, resulting in no significant difference in explained variance across all visual field maps (F_(1,185)_=0.59, *n.s.*).

##### Definition of V1-VO1

Maps of pRF phase and eccentricity were projected onto an inflated cortical surface reconstruction for each subject (**Figs S3–S6**). Borders between retinotopic maps were drawn on the cortical surface down the center of phase transitions occurring at the vertical or horizontal meridian^16,60,61^. V1, hV4^61,62^, and VO1^61,62^ were drawn as hemifields representing the contralateral visual field. V2 and V3 were drawn as quarterfields separated by V1, and were later combined to produce a hemifield representation. Individual maps were drawn by JG and independently checked by VN and KGS.

##### Definition of face- and character-selective functional regions of interest (ROIs)

Statistical contrasts of faces or characters > all other stimuli were thresholded at t-values > 3 for all subjects, as in our previous work^23,27,56^. Face-selective voxels that responded more strongly to faces than other stimuli and were located in the posterior lateral fusiform gyrus were defined as pFus-faces/FFA1. Character-selective voxels that responded more strongly to pseudowords and number strings than all other stimuli that were located on the posterior occipitotemporal sulcus lateral to pFus-faces were defined as pOTS-chars as in^56^. This region is also defined elsewhere as VWFA1^56,63^ using real word stimuli. Given that our region (pOTS-chars) occupies the same anatomical location as VWFA1, we refer to it throughout the manuscript as word-selective cortex for simplicity.

##### Estimating population receptive fields (pRF)

After functional data were transformed to the whole brain anatomy and restricted to the cortical ribbon, a population receptive field model was fit in each voxel^17^. For each voxel, a 2-dimensional Gaussian receptive field is modeled, having a center described by x and y coordinates and a sigma describing the width, and a parameter, g, describing its gain. An additional variable is fit for each voxel describing a compressive summation factor of the product of the stimulus and the Gaussian receptive field to better describe nonlinear summation properties of cortical responses as one ascends the visual hierarchy^36^. A candidate timecourse is produced from this pRF by convolving an HRF with the product of the stimulus movie and the pRF. The variables x, y, and sigma are swept until the variance explained of the voxel’s timecourse is maximized by the pRF model. Voxels were only included for subsequent analysis if the variance explained by the pRF model was greater than 5%. Additionally, to ensure the most accurate pRF fits, voxels whose pRF centers were outside the stimulus field (>7° radial eccentricity) or whose sigma was assigned the model’s minimum/floor value (0.21°) were excluded from further analysis.

##### pRF size versus eccentricity fits (**Fig 2**)

To evaluate the relationship between a pRF’s size and its eccentricity, voxels within an individual’s ROI were entered into a linear regression in which each voxel’s contribution was weighted by the variance explained of the pRF model. Only voxels with greater than 5% variance explained were included. The line-of-best fit was derived in each subject for each ROI, and then the slope and intercept of this line was averaged across participants of each age group.

##### Visual field coverage analyses

To calculate the visual field coverage (VFC) for a given ROI and subject, all voxels in an ROI that contain pRFs with >5% variance explained by the model are included and modeled as a Gaussian with a peak normalized to 1. The VFC is produced at each point by averaging the value across pRFs that cover that point, and then normalizing by the maximum coverage value in that subject. We also implemented a bootstrapping procedure^59^ that draws with replacement n-voxels from a subject’s ROI of size n, and produces an average VFC from 50 iterations to reduce the effect of outlier voxels. The average VFC from this bootstrapping approach is the VFC used for a given subject’s ROI. To produce the average VFC of subjects in each age group (**Figs 2** and **4**), the VFC is averaged across subjects of an age group. For the VFC of the visual field maps shown in **Fig 2** we first flipped for each subject the VFC of right hemisphere map over the vertical meridian and averaged with left hemisphere VFC before averaging across subjects. To measure the extent of the VFC for face- and word-selective regions (**Fig 4F),** we estimated the bilateral VFC for pFus-faces and pOTS-chars in each subject. pRF coverage density was binarized in each subject’s ROI (nonzero coverage assigned a value of 1) and the proportion of the visual field covered was multiplied by the total area stimulated by the sweeping bar stimulus (pr^2^, r=7^o^), resulting in the square degrees of visual angle covered by an individual’s ROI. We then averaged this across subjects in a group.

##### Center-of-mass distance from fixation

To quantify the foveal bias observed in face- and character-selective regions, we computed the center-of-mass (CoM) distance of the VFC of each region from the center of the visual field (**Fig 4**). This was derived by multiplying each coordinate by the normalized coverage density to obtain the center of VFC in a given region within children or adults. This measure was then jackknifed, repeated n times leaving out n-1 subjects on each fold, to produce the bars of standard error.

##### Measuring fixation patterns during free viewing of face and word stimuli outside the scanner

All subjects completed a recognition memory behavioral experiment while being eye-tracked with an Eyelink 1000 eyetracker (www.sr-research.com) in our eye tracking lab before scanning. Participants were seated, head-fixed using a chin rest and positioned 54cm from a monitor and told to freely view stimuli. The experiment had 3 parts: (1) *Encoding:* participants viewed images from five categories (child/adult faces, indoor/outdoor scenes, car/guitar objects, word/number characters, whole/limb bodies) and performed a 1-back task, indicating when 2 consecutive images were identical. Visual stimuli subtended 4° to 7° of visual angle, presented centrally within a 9^o^ square (see Fig 4 for examples). (2) *Fixation:* participants were instructed to fixate on a central dot while viewing a rotating checkerboard. This part was 4 minute long and served to train the participants to fixate during subsequent pRF mapping experiments. (3) *Recognition memory:* immediately followed the fixation training. Here, participants were presented with a surprise recognition task in which images appeared on the screen and for each image they were asked to indicate if it was previously seen during the “encoding” phase or if was a new image. This part was self-paced and the images appeared on the screen until participants made a decision. We report fixation from this phase.

##### Eye movement analysis

After removing timepoints during which participants blinked, data presented in **Fig 5** were analyzed in the following way: Fixation patterns were plotted in a 2-dimensional matrix (768x1024 pixel grid, equal in size to the stimulus presentation screen) and smoothed with a small Gaussian filter (sigma=18.75pixels) for the purpose of averaging data across subjects. Fixation density was normalized by the maximum in each subject, and then averaged for a given stimulus across all participants of an age group. The adult average fixation density was thresholded at 70% overlap for each stimulus and defined as the “adult fixation zone“, AFZ. The ratio of individual fixations made inside versus outside this this AFZ was calculated for each child participant and image, and then averaged across participants and stimuli of a given class (e.g. faces). The ratio was defined as (fixation time outside AFZ) / (total fixation time). A value of 1 indicates that all fixations occurred outside the AFZ, and value of 0 indicates that all fixations were within the AFZ. We then calculated if children fixated outside the AFZ significantly higher than chance, chance here being that 30% of fixations would occur outside the AFZ (as it was defined in adults as the 70% overlap contour).

##### Fixation bias vector analysis

The average fixation density for each face and word stimulus from the visual recognition test was calculated separately for children and adults. We first calculated for each image the center-of-mass of the distribution of adult fixations, similarly to the adult-fixation zone analysis discussed above, finding the center of the zone where 70% of adults fixated. We then calculated the center of mass of child fixations. From this center, a vector was produced pointing towards the center of fixation density on the same stimulus in children. Bias in child fixation vectors (**Fig 5C**) was quantified using a t-test to determine if vectors for a given stimulus category significantly deviated away from a vector bisecting the quadrant where the visual field coverage exists (for example, the coverage of right pFus-faces in the lower left quadrant) which we term the null quadrant. This procedure tested the hypothesis that children fixate in an optimal manner (e.g., they do not fixate in such a way that would move their limited coverage away from the informative region in the stimulus). It was assumed that 25% of randomly distributed vectors would have angles within the null quadrant (if randomly distributed, 25% of vectors should lie in quadrant spanning a quarter of the visual field), and t-tests were performed to assess if resulting bias vectors significantly deviated from this null.

##### Statistical Analysis

N-way ANOVAs were run for data presented in **Fig 2** with appropriate grouping variables and revealed no main effects or interactions, and thus no t-tests or KS-tests were performed. For **Fig 3**, N-way ANOVAs were run for pRF size and eccentricity treating ROI, hemisphere, and age groups as separate variables. For all statistical tests we report any significant main effects or interactions. Data going into ANOVAs was tested for normality assumptions using a Lilliefors test, and all data met or were very close to normal. ANOVAs are robust against modest deviations from normality, and no data populations have any gross violation of normality (as visible in the bar plots). All t- or KS-tests conducted were two-tailed. To test if any of our effects were correlated with age, we calculated the Pearson correlation coefficient on data underlying **Figs 2** and **4**; These values are reported in the text with the number of subjects going into each correlation. None of these correlations were significant. Bootstrapping methods were used to produce VFC plots in **Fig 2** and **4** to ensure robustness of fits and downweight outlier voxels; this bootstrapping method is described in the section <u>Visual field coverage analyses</u>, above. For **Fig 4A-D**, a two dimensional, 2-sample Kolmorogov-Smirnov test^64^, which is a nonparametric test comparing two continuous distributions simultaneously along two dimensions, was run on each ROI to test if the VFC was different across age-groups. All errorbars in the main and supplementary figures represent standard error of the mean across subjects.

##### Data and Code Availability

All data will be made available upon request. Analysis code is on github.com/VPNL and available upon request.

## AUTHOR CONTRIBUTION

J.G. contributed to experimental design, collection and analysis of data, and preparation of the manuscript. B.J., M.B., and V.N. contributed to collection and analysis of data, and preparation of the manuscript. K.G.-S. contributed to the experimental design, analysis of data, and manuscript preparation.

**Supplementary Figure 1:**
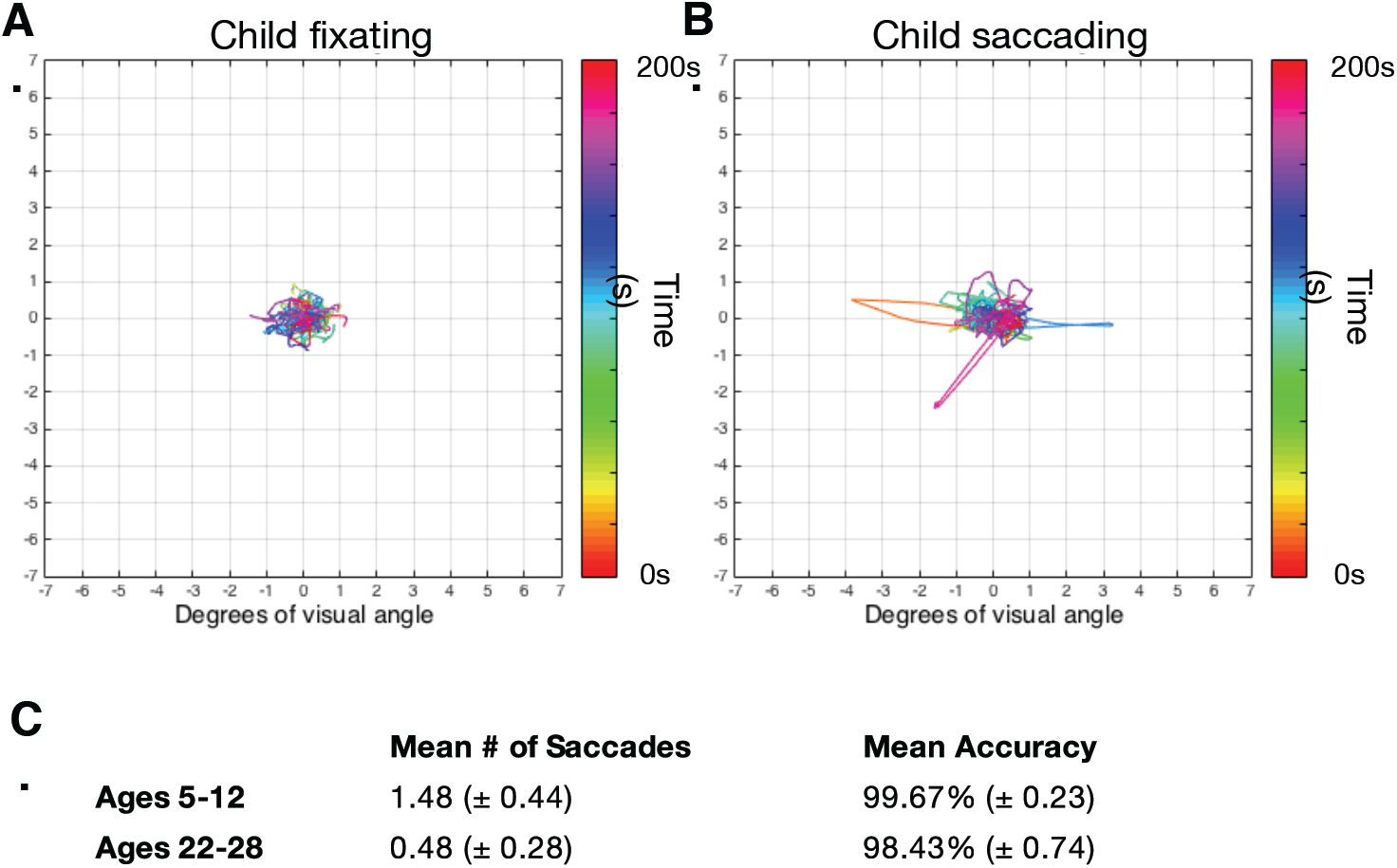
Fixation and behavioral performance during retinotopic scanning. Fixation patterns from example subjects either fixating(A) or (B) making minor saccades. The fixation path is color coded according to time (seconds) during the retinotopic mapping. Small deviations from the center are likely microsaccades and pupil-tracking noise from the scanner environment. (C) Behavioral performance during pRF mapping. Numbers indicate mean and standard deviation.

**Supplementary Figure 2:**
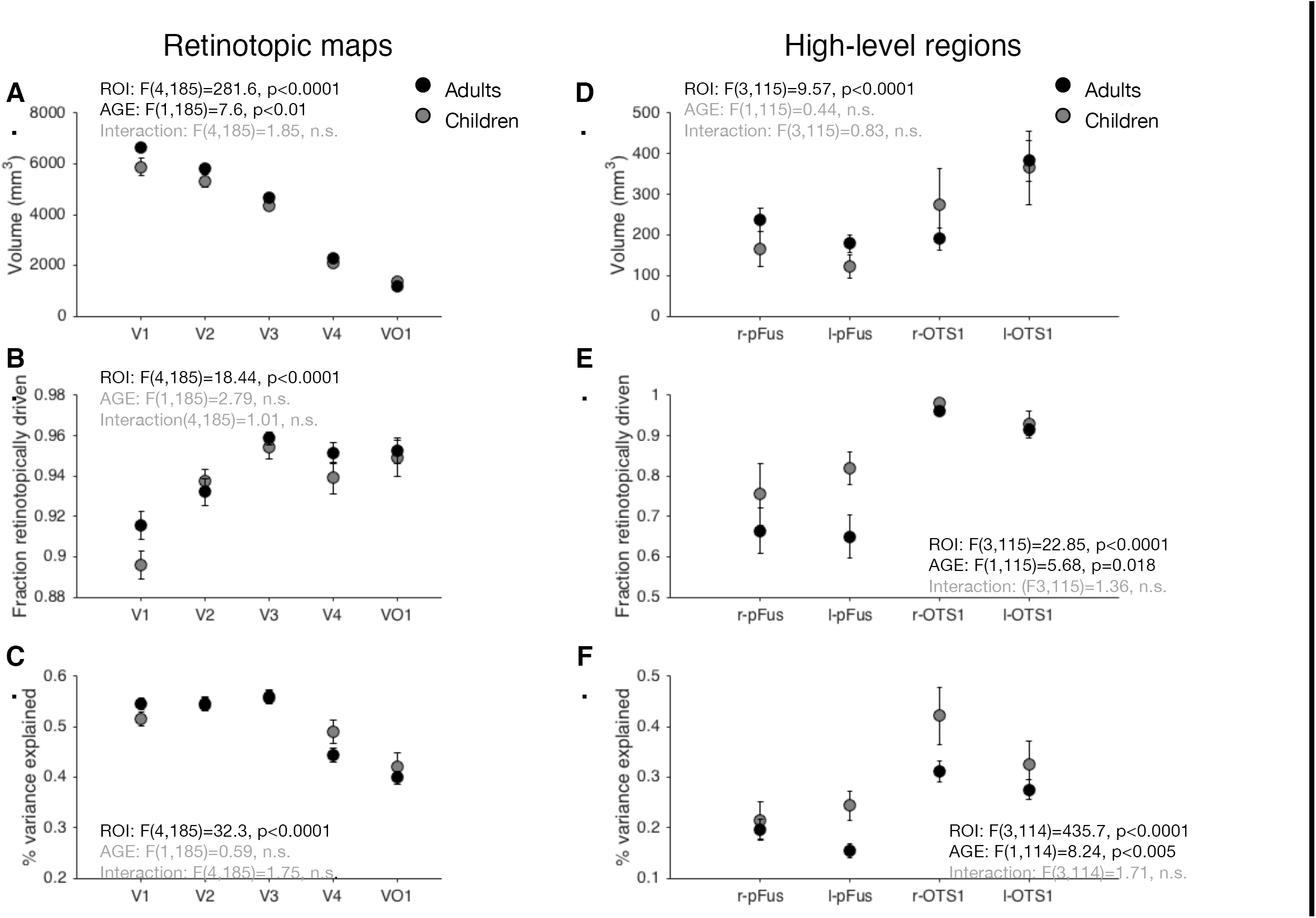
Descriptive data of retinotopic maps V1-VO2 and regions of VTC. **(A)** Volume in cubic millimeters is reported from visual field maps V1 through VO1. Error bars represent standard error. Reported volume measurements are the volume of voxels within the map that survive variance-explained thresholding. Children are light gray, adults black. Subjects included are matched for variance explained in V1. Subject numbers are the same as those reported in Figures 1 and 4. **(B)** The proportion of an ROI that is retinotopically driven above the 5% variance-explained threshold. Children are gray, adults black. **(C)** The mean percentage of variance explained across ROIs in children (gray) and adults (black) after variance-explained thresholding. **(D-F)** Same as A-C but for face- and word-selective regions. The r- and l-denote right and left hemisphere. ANOVAs for each were run with grouping variables of ROI and age-group. Main factors are reported, followed by the interaction.

**Supplementary Figure 3:**
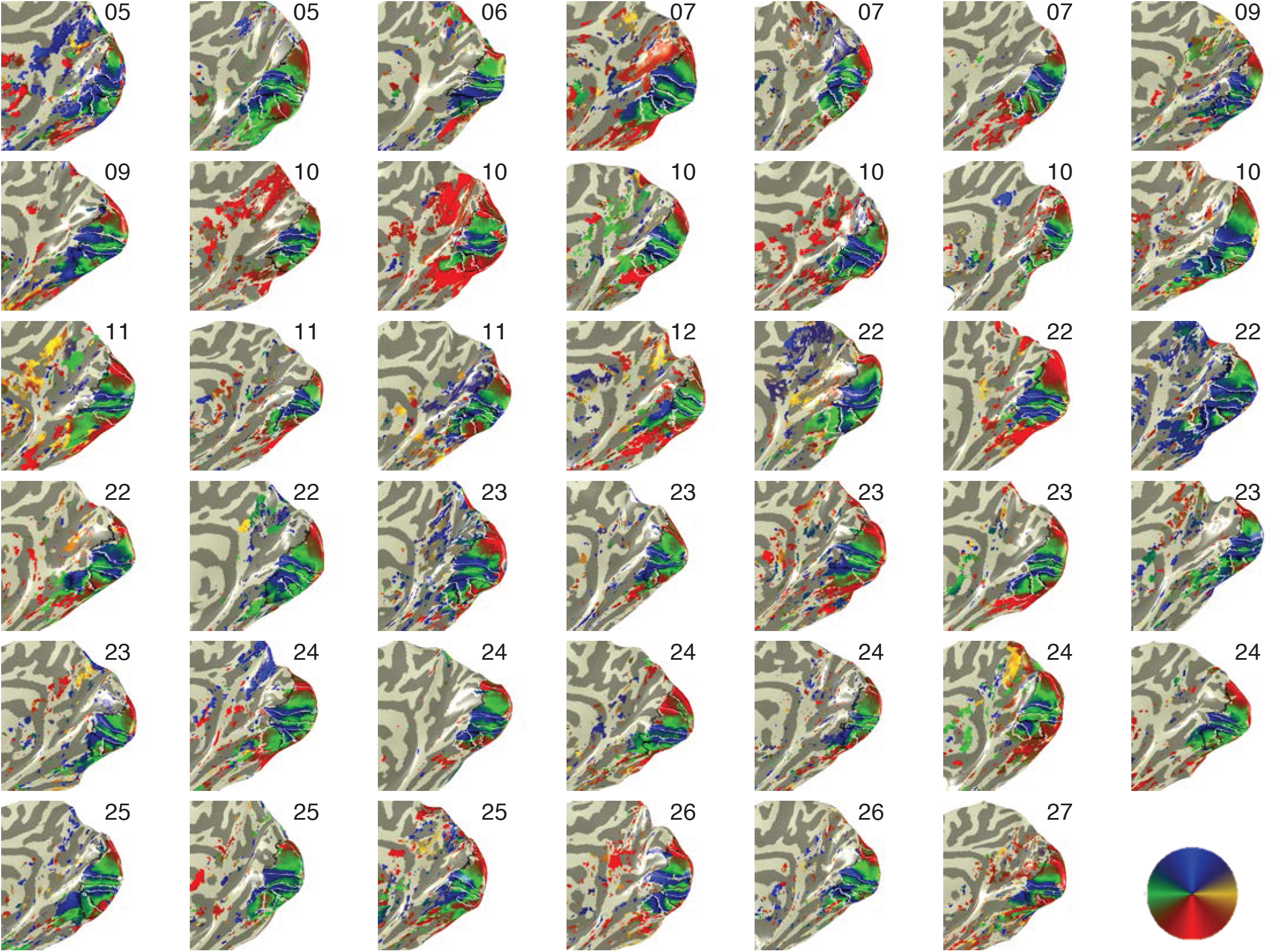
Polar angle maps of the right hemisphere occipital and temporal lobes for all subjects. Voxels are thresholded at 5% variance explained. All maps that we defined are presented, including V1, V2, V3, hV4, VO1. Not all maps could be delineated in each subject. Numbers indicate the age of the participant. *Color wheel:* polar angle color coding.

**Supplementary Figure 4:**
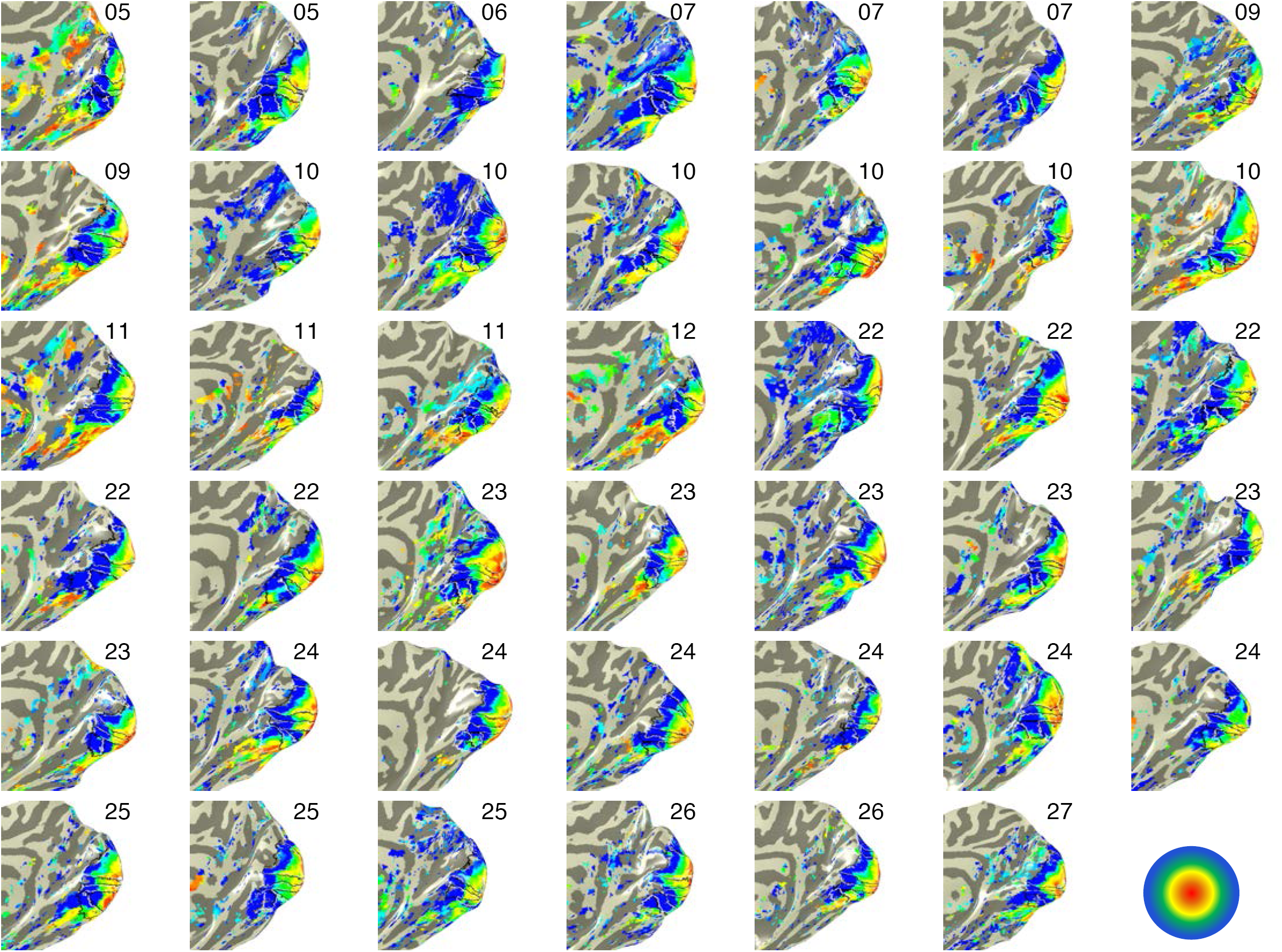
Eccentricity maps of the right hemisphere occipital and temporal lobes for all subjects. Voxels are thresholded at 5% variance explained. All maps that we defined are presented, including V1, V2, V3, hV4, VO1. Not all maps could be delineated in each subject. Numbers indicate the age of the participant. *Color wheel:* eccentricity color coding.

**Supplementary Figure 5:**
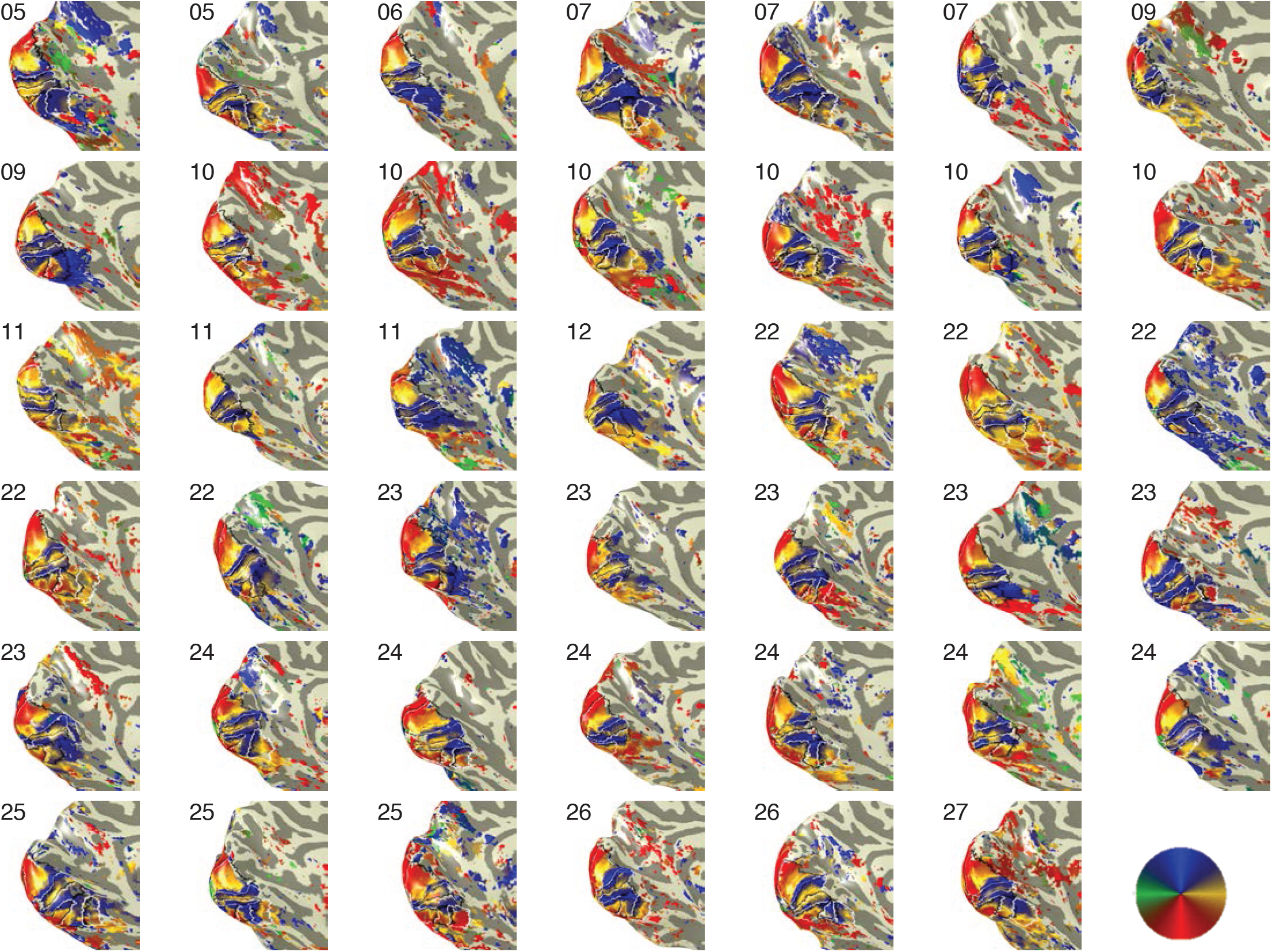
Polar angle maps of the left hemisphere occipital and temporal lobes for all subjects. Voxels are thresholded at 5% variance explained. All maps that we defined are presented, including V1, V2, V3, hV4, VO1. Not all maps could be delineated in each subject. Numbers indicate the age of the participant. *Color wheel:* polar angle color coding.

**Supplementary Figure 6:**
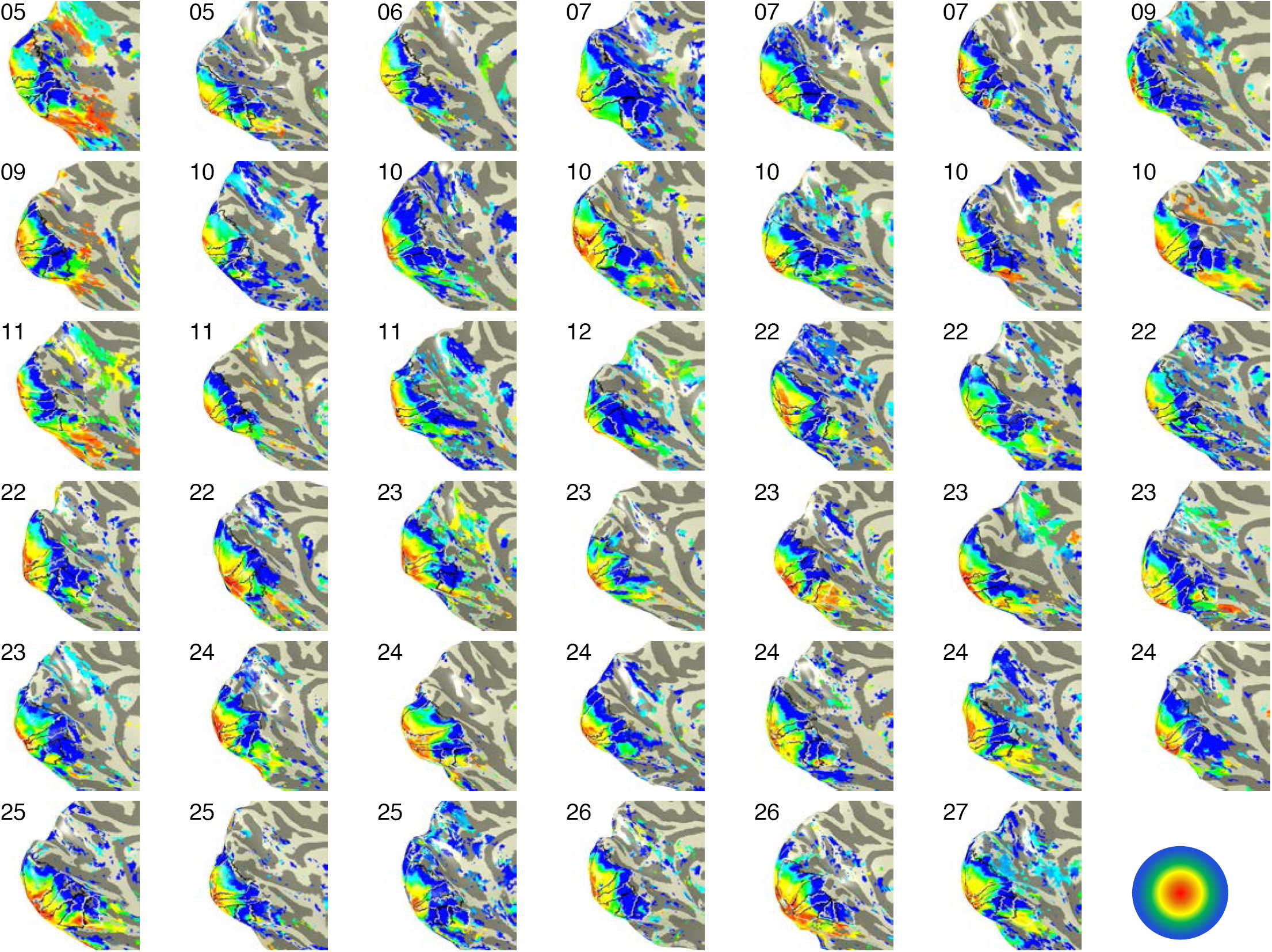
Eccentricity maps of the left hemisphere occipital and temporal lobes for all subjects. Voxels are thresholded at 5% variance explained. All maps that we defined are presented, including V1, V2, V3, hV4, VO1. Not all maps could be delineated in each subject. Numbers indicate the age of the participant. *Color wheel:* eccentricity color coding.

**Supplementary Figure 7:**
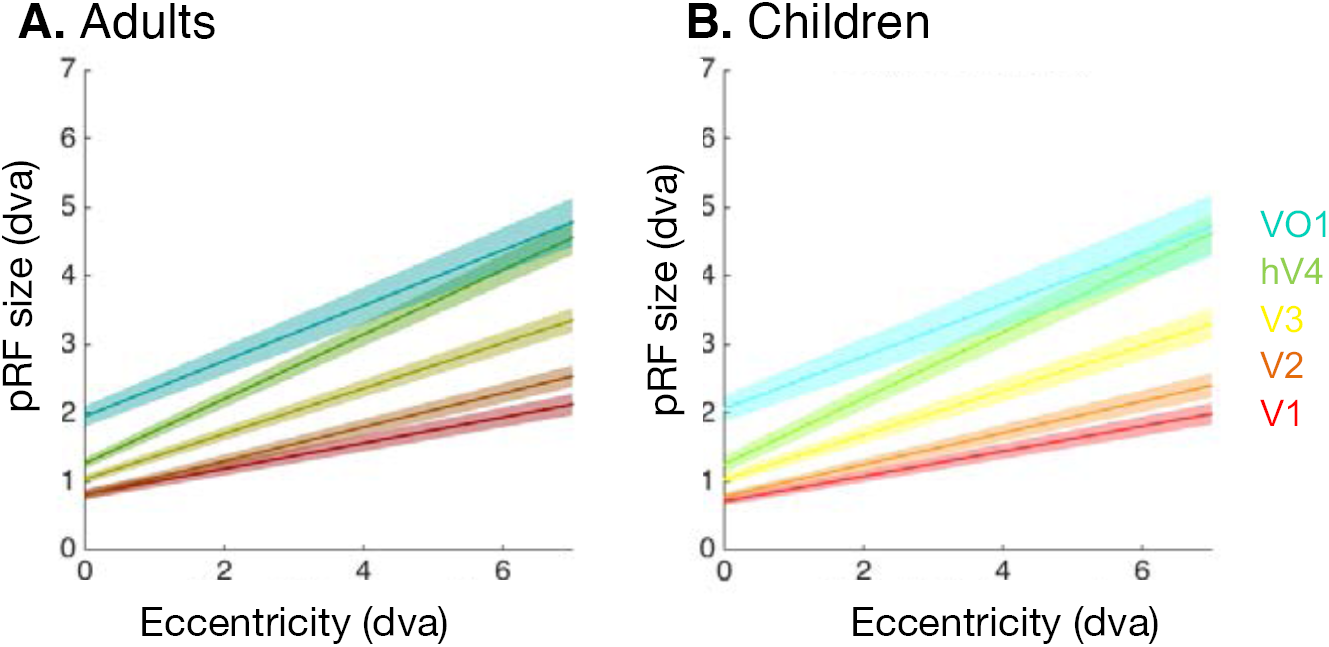
pRF size versus eccentricity fits in children and adults are similar. The line of best fit (solid line) and the standard error (shaded region) illustrates the relationship between pRF eccentricity and size in units of degrees of visual angle (dva). **(A)** Fits for V1 through VO1 are plotted for 23 adults. **(B)** Same as A but for 18 children.

**Supplementary Figure 8:**
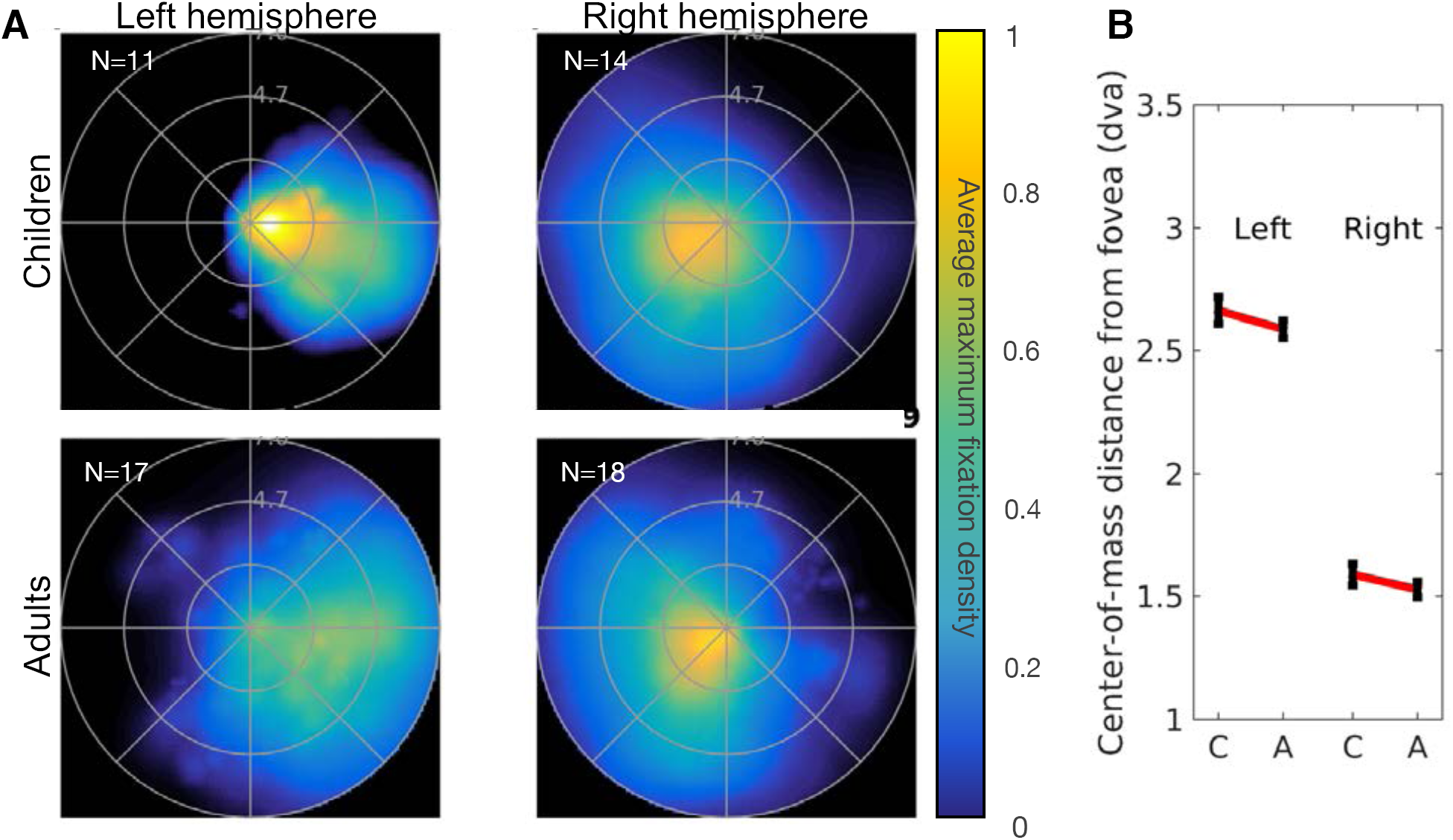
Visual field coverage in matched size pFus-faces. Left and right pFus-faces in children were dilated to match the average adult ROI size. (A) Average visual field coverage in children (top) and adults (bottom) In the left hemisphere the visual field coverage became less foveal from childhood to adulthood, and in the right hemisphere the visual field coverage became more foveal. (B) The center of mass (CoM) of the visual field coverage. *Error bars:* SEM. Results are similar to main data presented in Figure 4, suggesting ROI size is not a factor driving the effect of pRF coverage development.

**Supplementary Figure 9:**
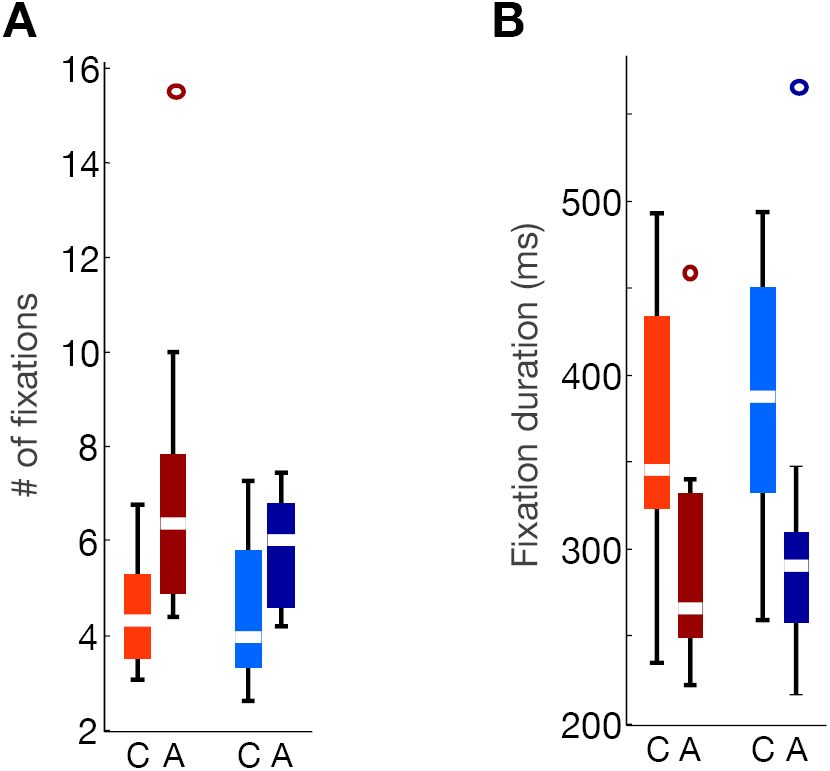
Fixation number and duration on face and pseudoword stimuli in children and adults during the recognition task. (A) The of number of fixations on faces and pseudowords in children (light colors) and adults (darker color) during the recognition task. Adults make significantly more fixations than children (t(21)=2.7, p<0.02), but they are clustered near the center of the stimulus (**Fig 5**). (B) Fixation duration on faces and pseudowords in children (light colors) and adults (dark colors) during the recognition task. Adults’ fixation durations are significantly shorter compared to those of children (t(21)=2.4, p<0.03). *White:* median; *Box:* 25^th^ and 75^th^ percentiles; *Whiskers:* range; *Circles:* outliers; *C:* children; *A:* adults; *red:* unfamiliar faces; *blue:* pseudowords.

## REFERENCES

1. Wilmer, T. Discharge Patterns and Functional of Mammalian Retina *. J Neurophysiol. 16, 37–68, (1951).

2. Hubel, D. H. & Wiesel, T. N. Receptive fields and functional architecture in two nonstriate visual areas (18 And 19) of the cat. J Neurophysiol 28, 229–289 (1965).

3. Arcaro, M. J., McMains, S. A., Singer, B. D. & Kastner, S. Retinotopic organization of human ventral visual cortex. J. Neurosci. 29, 10638–10652 (2009).

4. Kay, K. N., Weiner, K. S. & Grill-Spector, K. Attention reduces spatial uncertainty in human ventral temporal cortex. Curr. Biol. 25, 595–600 (2015).

5. Silson, E. H., Groen II, Kravitz, D. J. & Baker, C. I. Evaluating the correspondence between face-, scene-, and object-selectivity and retinotopic organization within lateral occipitotemporal cortex. J. Vis. 16, 14 (2016).

6. Le, R., Witthoft, N., Ben-Shachar, M. & Wandell, B. The field of view available to the cortical reading circuitry. bioRxiv (2016). doi: https://doi.org/10.1101/069369

7. Drover, J. R., Morale, S. E., Wang, Y.-Z., Stager, D. R. & Birch, E. E. Vernier acuity cards: examination of development and screening validity. Optom. Vis. Sci. 87, E806–E812 (2010).

8. Weigelt, S. et al. Domain-specific development of face memory but not face perception. Dev. Sci. 17, 47–58 (2014).

9. Dalton, K. M. et al. Gaze fixation and the neural circuitry of face processing in autism. Nat Neurosci 8, 519–526 (2005).

10. Rayner, K. Eye movements in Reading and Information Processing: 20 Years of Research. Psychol. Bull. 124, 372–422 (1998).

11. Ungerleider, L. G. & Mishkin, M. in Analysis of visual behavior (eds. Ingle, D. J., Goodale, M. A. & Mansfield, R. J. W.) 549–586 (MIT Press, 1982).

12. Kanwisher, N., McDermott, J. & Chun, M. M. The fusiform face area: a module in human extrastriate cortex specialized for face perception. J Neurosci 17, 4302–4311 (1997).

13. Cohen, L. et al. The visual word form area: spatial and temporal characterization of an initial stage of reading in normal subjects and posterior split-brain patients. Brain 123 (Pt 2, 291–307 (2000).

14. Parvizi, J. et al. Electrical stimulation of human fusiform face-selective regions distorts face perception. J. Neurosci. 32, 14915–14920 (2012).

15. Hirshorn, E. a. et al. Decoding and disrupting left midfusiform gyrus activity during word reading. Proc. Natl. Acad. Sci. U. S. A. 113, 201604126 (2016).

16. Wandell, B. A. & Winawer, J. Computational neuroimaging and population receptive fields. Trends Cogn. Sci. 19, 349–357 (2015).

17. Dumoulin, S. O. & Wandell, B. A. Population receptive field estimates in human visual cortex. Neuroimage 39, 647–660 (2008).

18. Malach, R., Levy, I. & Hasson, U. The topography of high-order human object areas. Trends Cogn Sci 6, 176–184 (2002).

19. Huberman, A. D., Feller, M. B. & Chapman, B. Mechanisms Underlying Development of Visual Maps and Receptive Fields. Annu. Rev. Neurosci. 31, 479–509 (2008).

20. White, L. E. & Fitzpatrick, D. Vision and cortical map development. Neuron 56, 327–38 (2007).

21. Luo, L. & Flanagan, J. G. Development of continuous and discrete neural maps. Neuron 56, 284–300 (2007).

22. Golarai, G. et al. Differential development of high-level visual cortex correlates with category-specific recognition memory. Nat Neurosci 10, 512–522 (2007).

23. Natu, V. S. et al. Development of Neural Sensitivity to Face Identity Correlates with Perceptual Discriminability. J. Neurosci. 36, 10893–10907 (2016).

24. Scherf, K. S., Behrmann, M., Humphreys, K. & Luna, B. Visual category-selectivity for faces, places and objects emerges along different developmental trajectories. Dev. Sci. 10, F15–30 (2007).

25. Cantlon, J. F., Pinel, P., Dehaene, S. & Pelphrey, K. A. Cortical representations of symbols, objects, and faces are pruned back during early childhood. Cereb. Cortex 21, 191–199 (2011).

26. Peelen, M. V., Glaser, B., Vuilleumier, P. & Eliez, S. Differential development of selectivity for faces and bodies in the fusiform gyrus. Dev. Sci. 12, (2009).

27. Gomez, J. et al. Microstructural proliferation in human cortex is coupled with the development of face processing. Science (80-.). 355, 68–71 (2017).

28. Sowell, E. R. et al. Mapping cortical change across the human life span. Nat. Neurosci. 6, 309–315 (2003).

29. Conner, I. P., Sharma, S., Lemieux, S. K. & Mendola, J. D. Retinotopic organization in children measured with fMRI. J. Vis. 4, 509–523 (2004).

30. Behrmann, M. & Plaut, D. C. A vision of graded hemispheric specialization. (2015).

31. Dehaene, S., Cohen, L., Morais, J. & Kolinsky, R. Illiterate to literate: behavioural and cerebral changes induced by reading acquisition. Nat. Rev. Neurosci. 16, 234–244 (2015).

32. Dehaene, S. et al. How learning to read changes the cortical networks for vision and language. Science (80-.). 330, 1359–1364 (2010).

33. Yarbus, A. L. in Eye movements and vision (ed. Yarbus, A. L.) 171–211 (Springer, 1967).

34. Schyns, P. G., Bonnar, L. & Gosselin, F. Show me the features! Understanding recognition from the use of visual information. Psychol. Sci. 13, 402–409 (2002).

35. Zhang, L., Tong, M. H., Marks, T. K., Shan, H. & Cottrell, G. W. SUN: A Bayesian framework for saliency using natural statistics. J. Vis. 8, 32.1–20 (2008).

36. Kay, K. N., Winawer, J., Mezer, A. & Wandell, B. A. Compressive spatial summation in human visual cortex. J. Neurophysiol. 110, 481–494 (2013).

37. Silver, M. A. & Kastner, S. Topographic maps in human frontal and parietal cortex. Trends in Cognitive Sciences 13, 488–495 (2009).

38. Dundas, E. M., Plaut, D. C. & Behrmann, M. An ERP investigation of the co-development of hemispheric lateralization of face and word recognition. Neuropsychologia 61, 315–323 (2014).

39. Sacchi, E. & Laszlo, S. An event-related potential study of the relationship between N170 lateralization and phonological awareness in developing readers. Neuropsychologia 91, 415–425 (2016).

40. Maurer, U., Rossion, B. & McCandliss, B. D. Category specificity in early perception: face and word n170 responses differ in both lateralization and habituation properties. Front. Hum. Neurosci. 2, 18 (2008).

41. Rossion, B. Constraining the cortical face network by neuroimaging studies of acquired prosopagnosia. Neuroimage 40, 423–426 (2008).

42. Fox, C. J., Iaria, G. & Barton, J. J. Disconnection in prosopagnosia and face processing. Cortex. 44, 996–1009 (2008).

43. Behrmann, M. & Plaut, D. C. A vision of graded hemispheric specialization. Ann. N. Y. Acad. Sci. 1359, 30–46 (2015).

44. Moerel, M., De Martino, F. & Formisano, E. Processing of Natural Sounds in Human Auditory Cortex: Tonotopy, Spectral Tuning, and Relation to Voice Sensitivity. J. Neurosci. 32, 14205–14216 (2012).

45. Barton, B., Venezia, J. H., Saberi, K., Hickok, G. & Brewer, A. A. Orthogonal acoustic dimensions define auditory field maps in human cortex. Proc. Natl. Acad. Sci. U. S. A. 109, 20738–20743 (2012).

46. Huang, R.-S., Chen, C., Tran, A. T., Holstein, K. L. & Sereno, M. I. Mapping multisensory parietal face and body areas in humans. Proc. Natl. Acad. Sci. U. S. A. 109, 18114–9 (2012).

47. Harvey, B. M., Klein, B. P., Petridou, N. & Dumoulin, S. O. Topographic representation of numerosity in the human parietal cortex. Science (80-.). 341, 1123–1126 (2013).

48. Kersey, A. J. & Cantlon, J. F. Neural tuning to numerosity relates to perceptual tuning in 3- to 6-year-old children. J. Neurosci. 37, 512–522 (2016).

49. Duchaine, B. C. & Nakayama, K. Developmental prosopagnosia: a window to content-specific face processing. Curr. Opin. Neurobiol. 16, 166–173 (2006).

50. Schwarzkopf, D. S., Anderson, E. J., de Haas, B., White, S. J. & Rees, G. Larger Extrastriate Population Receptive Fields in Autism Spectrum Disorders. J. Neurosci. 34, 2713–2724 (2014).

51. Grill-Spector, K., Golarai, G. & Gabrieli, J. Developmental neuroimaging of the human ventral visual cortex. Trends Cogn Sci 12, 152–162 (2008).

52. Golarai, G., Liberman, A., Yoon, J. M. & Grill-Spector, K. Differential development of the ventral visual cortex extends through adolescence. Front. Hum. Neurosci. 3, 80 (2010).

53. Yeatman, J. D., Wandell, B. A. & Mezer, A. A. Lifespan maturation and degeneration of human brain white matter. Nat. Commun. 5, 4932 (2014).

54. Mezer, A. et al. Quantifying the local tissue volume and composition in individual brains with magnetic resonance imaging. Nat. Med. 19, 1667–1672 (2013).

55. Feinberg, D. A. & Setsompop, K. Ultra-fast MRI of the human brain with simultaneous multi-slice imaging. J Magn Reson 229, 90–100 (2013).

56. Stigliani, A., Weiner, K. S. & Grill-Spector, K. emporal Processing Capacity in High-Level Visual Cortex Is Domain Specific. J. Neurosci. 35, 12412–12424 (2015).

57. Weiner, K. S. & Grill-Spector, K. Not one extrastriate body area: Using anatomical landmarks, hMT+, and visual field maps to parcellate limb-selective activations in human lateral occipitotemporal cortex. Neuroimage 56, 2183–2199 (2011).

58. Hummer, A. et al. Eyetracker-based gaze correction for robust mapping of population receptive fields. Neuroimage 142, 211–224 (2016).

59. Winawer, J. & Witthoft, N. Human V4 and ventral occipital retinotopic maps. Vis. Neurosci. 32, E020 (2015).

60. Sereno, M. I. et al. Borders of multiple visual areas in humans revealed by functional magnetic resonance imaging. Science (80-.). 268, 889–893 (1995).

61. Witthoft, N. et al. Where is human V4? Predicting the location of hV4 and VO1 from cortical folding. Cereb. Cortex 24, 2401–2408 (2014).

62. Brewer, A. A., Liu, J., Wade, A. R. & Wandell, B. A. Visual field maps and stimulus selectivity in human ventral occipital cortex. Nat Neurosci 8, 1102–1109 (2005).

63. Ben-Shachar, M., Dougherty, R. F., Deutsch, G. K. & Wandell, B. A. Differential sensitivity to words and shapes in ventral occipito-temporal cortex. Cereb. Cortex 17, 1604–1611 (2007).

64. Fasano, G. & Franceschini, A. A multidimensional version of the Kolmogorov-Smirnov test. Mon. Not. R. Astron. Soc. 155–170 (1987).

